# ‘Wanting’ versus ‘Needing’ related value: an fMRI meta-analysis

**DOI:** 10.1101/2021.09.24.461758

**Authors:** J Bosulu, M.-A. Allaire, L. Tremblay-Grénier, Y. Luo, S. Eickhoff, S. Hétu

## Abstract

Consumption and its excesses are sometimes explained by imbalance of need or lack of control over ‘wanting’. ‘Wanting’ assigns value to cues that predict rewards, whereas ‘needing’ assigns value to biologically significant stimuli that one is deprived of. Here we aimed at studying how the brain activation patterns related to value of ‘wanted’ stimuli differs from that of ‘needed’ stimuli using ALE neuroimaging meta-analysis approaches. We used the perception of a cue predicting a reward for ‘wanting’ related value and the perception of food stimuli in a hungry state as a model for ‘needing’ related value. We carried out separate, contrasts, and conjunction meta-analyses to identify differences and similarities between ‘wanting’ and ‘needing’ values. Our overall results for ‘wanting’ related value show consistent activation of the ventral tegmental area, striatum and pallidum, regions that both activate behaviour and direct choice; while for ‘needing’ related value we found an overall consistent activation of the middle insula and to some extent the caudal-ventral putamen, regions that only direct choice. Our study suggests that wanting has more control on consumption, and a needed stimuli must become wanted in order to be pursued.

## INTRODUCTION

Current consumption (e.g. food, transport, etc.) in western countries seems to be one of the causes of ecological problems we are facing (Lipschutz, 2001). According to some, this consumption is in part due to the fact that we consume what we want beyond what we need (Stearn, 2006). Apart from problems related to ecology, in our daily behaviors such as those related to food consumption, excesses and maladaptive behaviors are sometimes explained by an imbalance of need or a lack of control over ‘wanting’. Indeed, Campbell (1998) reports that research on consumer behavior has shown that there are two types of rhetoric used to justify the action of purchase: needs and wants, as well as their synonyms. He also argues that rhetoric of needs is derived from utilitarianism and puritanism that advocated a life based on necessity or satisfaction; while the rhetoric of wants is based on romanticism and linked to the pursuit of pleasure (Campbell, 1998). Beyond this rhetorical distinction, one can wonder: is there a true difference between something that we need and something that we want? At the neural level, needing is related to a state of deprivation of something important for life or survival (Bouton, 2016), and increases arousal through interoceptive salience (Craig, 2003). ‘Wanting’ is related to the prediction of reward in the brain and is more closely related to motivation (Berridge, 2004). Although earlier theories suggested that need or deprivation defines motivation (Hull, 1943), it was later demonstrated that cues that signal hunger do not elicit motivation to eat, while food-related cues did lead to motivation to eat (Bindra, 1974), showing that motivated behaviours are more determined by reward prediction (which is closer to ‘wanting’) than need state, though the later can have multiplicative effect (Toates, 1994). The distinction might seem clear. However, since ‘needed’ stimuli are often pursued and thus associated with motivational value, it’s not obvious whether an external stimulus, such as food, is pursued because of its ‘needing’ or ‘wanting’ related value. As ‘needed’ stimuli might have a different form of value than ‘wanted’ stimuli, we can wonder how in the brain the value of ‘needed’ stimuli differs from that of ‘wanted’ stimuli.

Given that we cannot test wanting and/or needing as general phenomena, here we are testing some manifestation of them. Hence, to conceptualize this distinction, we refer to “wanting a stimulus” as ‘Wanting’_ST_, which represents a brain reaction to a reward predicting cue that triggers reward seeking. We refer to “needing a stimulus” as ‘Needing’_ST_, which represents the brain reaction to a stimulus that one is deprived of, without necessarily seeking it. We propose to use activation pattern to the perception of a cue predicting a reward as a model for ‘Wanting’_ST_ related value; and to use the activation pattern during the perception of a food stimulus while in the state of hunger as a model for ‘Needing’_ST_ related value (Silverman et al., 2014; Spear, 2011). In order to answer our question, we used a neuroimaging meta-analytic approach, comparing the patterns of brain activations during the processing of ‘Wanting’_ST_ versus ‘Needing’_ST_. Previous meta-analyses have focused on either ‘Wanting’_ST_ (Wilson et al., 2018; Oldham et. al., 2018; Sescousse et. al. 2013) or ‘Needing’_ST_ (van der Lan et al., 2011; LaBar et. al., 2001; Chen & Zeffiro, 2020), but no work has directly compared both activation patterns.

### ‘Wanting’_ST_ related value

Non-human animals tend to respond and ‘want’ food even when they are no longer hungry (Bouton, 2016). This is also the case for humans: cues of M&M or pictures of cigarettes (for smokers) lead to more consumption even after having been consumed to satiety (Watson et al., 2014; Hogarth & Chase, 2011). ‘Wanting’_ST_ is a concept from incentive salience theory that comes from animal studies (Berridge & Robinson, 1998; see also: Ikemoto & Panksepp, 1999; Salamone & Correa, 2002; Salamone, et al., 1997) which states that ‘Wanting’_ST_ is based on two neuropsychological processes: the first is a pavlovian cue that predicts the reward; and the second is the dopaminergic state (which might be enhanced by hunger, thirst, emotions, drugs, etc.) (Berridge, 1996). In other words,‘Wanting’_ST_ depends on external stimuli that act as pavlovian cues that predict rewards (Berridge, 2018). The attribution of value to these cues depends on mesolimbic dopamine (Berridge, 1996). The latter is secreted within the ventral tegmental area (VTA) by a reward cue (Schultz, 1998), and projected within the nucleus accumbens (Nacc) and ventral pallidum (Warlow & Berridge, 2021; Zhang et al. 2009). In this sense, the VTA would be more related to change in reward prediction (Schultz, 2015; Schultz et. al., 1997) and its phasic activation determines directional value (preference/choice or action selection), while the NAcc is more related to value attribution (an incentive salience) to that reward prediction (Berridge and Aldrige, 2009; Lex and Hauber, 2008; Hamid et al., 2016), along with the central nucleus of the amygdala (Warlow & Berridge, 2021; Zhang et al. 2009; Balleine & Killcross, 2006). Thus, ‘Wanting’_ST_ starts with reward prediction. Human studies on reward prediction have shown the involvements of the orbitofrontal cortex (OFC) (O’Doherty et. al, 2002; O’Doherty, 2004), VTA (O’Doherty et. al, 2002; Krebs et. al., 2009; Carter et, al., 2009; Schott et. al., 2008; Oldham et. al., 2018), NAcc and ventral striatum (O’Doherty et. al, 2002; Knutson et. al 2003; O’Doherty, 2004; Knutson et. al 2001; Wilson et al., 2018; Carter et, al., 2009; Schott et. al., 2008; Oldham et. al., 2018; Simon et. al., 2015), amygdala (O’Doherty et. al, 2002; O’Doherty, 2004; Oldham et. al., 2018) and insula (O’Doherty et. al, 2002; Wilson et al., 2018; Oldham et. al., 2018). These results suggest that, in humans, the activation pattern of reward prediction which leads to ‘Wanting’_ST_ related value could implicate these regions.

### ‘Needing’_ST_ related value

A need state has the capacity to give and to control the preference/choice related value of a novel food or drink in relation to its consequence on the organism, once the organism has experienced the benefit of that stimulus in the need state (Dickinson & Balleine 1994; Balleine, 1992). Thus, ‘Needing’_ST_ related value can have an impact on choice and action selection (Dickinson & Balleine 1994) or preference (Salamone et al, 2018). For instance, hunger influences flavour preference learning in humans based on flavour (Brunstrom & Fletcher, 2008; Zellner et al., 1983), nutrients (Gibson et al., 1995; Kern et. al, 1993; Appleton et al., 2006), and odour-sweetness (Yeomans & Mobini, 2006). Moreover, the shifts in preference are found to persist beyond the initial training period (Brunstrom & Fletcher, 2008), suggesting a long term learned value. However, though ‘Needing’_ST_ provides directional value (preference/choice or action selection), in absence of reward prediction, ‘Needing’_ST_ (by itself) does not activate behavior (Bindra, 1974; Toates, 1994; Bolles and Moot, 1972; Berridge, 2004). In the brain, it has been suggested that ‘Needing’_ST_, which depends on interoception and its prediction and its prediction error (i.e., difference between predicted need state and actual need state) within the anterior insula and mid-posterior insula, respectively (Barrett & SImmon, 2015), also often recruits the anterior cingulate cortex (ACC) (Craig, 2003). Moreover, ‘Needing’_ST_ related value attribution implicates the OFC (Ostlund and Balleine, 2007; Balleine & O’Doherty, 2010), while ‘Needing’_ST_ related learning recruits the basolateral amygdala (Balleine and Killcross, 2006) and the long term association between external stimuli and their consequence on physiological need states recruits the caudate body and tail as well as the putamen, mostly the caudal-ventral putamen (Seger & Cincotta, 2005; Schwabe & Wolf, 2010; Kunimatsu et. al., 2019; Amita, et. al., 2018), and insular cortex (Balleine and Dickinson, 2000). Overall, previous fMRI meta-analyses and studies on hunger in humans have revealed regions associated with sensory integration, reward processing, and taste; including the insula (van der Lan et al., 2011; Goldstone, et. al., 2009; Siep et. al., 2009), the OFC (van der Lan et al., 2011; Goldstone, et. al., 2009; Siep et. al., 2009; Führer et. al., 2008), the amygdala (van der Laan, et. al., 2011; LaBar et. al., 2011; Führer, 2008; Mohanty, 2008; Goldstone, 2009), the dorsal striatum (van der Laan, et. al., 2011; Siep et. al., 2009), and the ACC (Goldstone, et. al., 2009; Siep et. al., 2009; Führer et. al., 2008); and many studies have found activations within the amygdala/parahippocampal gyrus (LaBar, et. a., 2011; Mohanty, et. al., 2008; Chen & Zeffiro, 2020). Hence hunger can cautiously be used as a proxy for ‘Needing’_ST_. Thus, based on the inherent association between hunger (and thirst) and ‘Needing’_ST_, the insula and ACC, and to some extent the OFC, amygdala, and caudate may be engaged in the processing of Needing’_ST_ related value and contribute to directional value.

### Two types of predictions and values

The conceptualization of ‘Wanting’_ST_ and ‘Needing’_ST_ here as processing of either wanted or needed stimuli implies two forms of predictions. Indeed, in both cases, the value of stimuli often depends on a prediction and a prediction error: in case of *‘*Wanting’_ST_, that prediction is related to reward (unexpected reward or reward predicting cue) and is computed in the ventral striatum (Takahashi et al., 2016), while the prediction error is computed in the VTA (Schultz, 2015). For ‘Needing’_ST_, that prediction is related to interoception (predicted state vs sensed state) and is said to be computed in visceromotor cortices (OFC, ACC, anterior insula), whereas the interoceptive prediction error is proposed to be computed within the mid-posterior insula (Barrett & Simmons, 2015). Moreover, both ‘Wanting’_ST_ and *‘*Needing’_ST_ establish a relation between the state (‘wanting’ state and *‘*needing’ state) and some external stimuli, where the state attributes some form of value to the stimuli. The value assigned to stimuli (by both ‘Wanting’_ST_ or *‘*Needing’_ST_) can have a directional effect or activational effect. The directional effect is linked to choice (preference or action selection) and directs towards or away from stimuli, while the activational effect is related to action and its initiation, maintenance, and vigor or effort (see Salamone et al., 2018). Indeed, *‘*Wanting’_ST_ depends on mesolimbic dopamine (Berridge, 1996) which provides both activational value (or effect) and directional value (or effect) to stimuli (see Salamone et. al., 2018). ‘Needing’_ST_ (by itself) doesn’t seem to provide the activational value that *‘*Wanting’_ST_ provides to stimuli (Berridge, 2004; see also Salamone et al., 2018). However, ‘Needing’_ST_ do provide directional value (Salamone et al., 2018; Balleine, 1992). Importantly, if both ‘Wanting’_ST_ and ‘Needing’_ST_ provide directional related value to stimuli that impacts choice, they do so in different ways. In the case of *‘*Wanting’_ST_, that choice value is pavlovian cue triggered and stimulus related (Berridge, 2012; Balleine, 2009). In the case of ‘Needing’_ST_, that choice value is act-outcome based (Balleine, 2009). Hence, depending on either state, stimulus value could be represented by different activation patterns in the brain (Dayan & Balleine, 2002; Berridge & Aldridge, 2009). However, no work has quantitatively tested this hypothesis.

As discussed, neuroimaging studies in the activation pattern of ‘Wanting’_ST_ related value shows consistent activation of the striatum, amygdala and insula (Knutson et. al 2003; O’Doherty, 2004; Knutson et. al 2001; O’Doherty et. al, 2002; Wilson et al., 2018; Oldham et. al., 2018; Carter et, al., 2009; Schott et. al., 2008). Those same regions have also been found in the activation pattern of ‘Needing’_ST_ (van der Lan et al., 2011; Goldstone, et. al., 2009; Siep et. al., 2009; LaBar et. al., 2011; Führer, 2008; Mohanty, 2008). It’s not clear how these regions contribute to either ‘Wanting’_ST_ or ‘Needing’_ST_. Our goal is thus to use a meta-analytic approach to compare the consistent brain activation patterns for ‘Wanting’_ST_ and ‘Needing’_ST_ related values by identifying similarities and differences between the brain activation patterns of these two states that guide value attribution and our consumption behaviors. To do this, we will quantitatively identify the consistent activation patterns after the observation of a reward cue/reward prediction (‘Wanting’_ST_), versus while (or after) observing a food cue when hungry (‘Needing’_ST_). We will then directly compare these activation patterns by conducting meta-analytic conjunction and contrast analyses.

## METHODOLOGY

We decided to use a meta-analytic approach as it provides an opportunity to quantitatively assess brain activation patterns of ‘Wanting’_ST_ vs. ‘Needing’_ST_ related values using large collections of data. This is useful as a summary of the existing literature is needed, not just because both concepts have rarely been directly compared and are often studied separately in neuroimaging studies, but also because each study might have low replicability, analytical and experimental flexibility, and/or small samples. Thus, our approach aims at identifying and comparing regions that are consistently activated for ‘Wanting’_ST_ and those that are consistently activated for ‘Needing’_ST_. Specifically, we first conducted two meta-analyses to quantitatively summarize results from functional magnetic resonance imaging (fMRI) published studies on the reward prediction for ‘Wanting’_ST_ (activation maps taken when participants received a reward predicting cue that triggers reward seeking); and on perceiving food stimulus while being hungry for ‘Needing’_ST_ (activation maps taken when participants perceived food while hungry). Second, we did a conjunction analysis to identify common regions that are consistently activated in both states. Finally, we contrasted ‘Wanting’_ST_ and ‘Needing’_ST_ consistent activation patterns by testing, [‘Wanting’_ST_-‘Needing’_ST_], and [‘Needing’_ST_-‘Wanting’_ST_].

### Included articles

Based on the view that ‘Wanting’_ST_ rests upon reward prediction that has been turned into a decision (see Berridge & Aldridge, 2009) we used the following keywords to identify articles related to ‘Wanting’_ST_:

> ((“prediction” AND “anticipation”) OR “desire” OR “wanting”)

Based on the view that ‘Needing’_ST_ such as hunger depends on interoception (Craig, 2003) coming from deprivation of something biologically important, we used the following keywords to identify articles related to ‘Needing’_ST_:

> (“alliesthesia” OR “interoceptive” OR “loss aversion” OR “need” OR “homeostasis” OR “modulating factor” OR “self-specificity” OR “self-referential” OR “hunger” OR “food deprivation”).

While the previous lists of keywords were specific to either ‘Wanting’_ST_ or ‘Needing’_ST_, the following keywords were the same for both ‘Wanting’_ST_ and ‘Needing’_ST_ ; those keywords were the following:

> (“reward” OR “motivation” OR “goal directed” OR “decision-making” OR “seeking” OR “incentive”) AND (“fMRI”)

These include words that are often conceptualized as related to ‘Wanting’_ST_ and to ‘Needing’_ST_ (Bouton, 2016; Panksepp, 2004).

For both ‘Wanting’_ST_ or ‘Needing’_ST,_, the following inclusion criteria were used: healthy subjects; whole-brain analyses (with or without SVC), MNI or Talairach Coordinates (all Talairach coordinates were converted to MNI SPM152 in Ginger ALE using Lacanster transform); maps were corrected (or cluster level corrected); activation contrast only.

With regard to ‘Wanting’_ST_, we typed the keywords on PubMed (February 2021). The database returned 159 articles. The main selection criteria were the presence of a cue that predicts a reward and triggers reward seeking contrasted with no prediction of reward (reward prediction>no reward prediction). After evaluation based on these criteria, 19 final articles were selected out of 26 that were fully read ; and from which we found three additional articles from reviews and other articles that met all the criteria for ‘Wanting’_ST_ for a total of 22 selected articles (see table 2 for list of retained articles). Note that these rewards were mostly money or points, so they are not (directly) related to food, but they are used because ‘Wanting’_ST_ or incentive motivation activates a general system, regardless of the type of stimulus (Bindra, 1968; Bouton, 2016). See Prisma in supplementary material for step by step exclusion of articles.

**Table 1.** Selection Criteria The red coloured “yes” means the criterion is crucial for the definition of either ‘Wanting’ST or ‘Needing’ST. Most mean BMI values were below 30, which is the threshold of obesity as defined by the World Health Organization (World Health Organization, 2006), however, we took people that had up to 30-35 of BMI, especially in contrasts when healthy and overweight were mixed. For the Millman, et al. (2019) study, we took the contrast of “large gain > small gain” (so exceptionally, we contrasted two rewards) ; first because there were no contrast for gain alone in general ; and because small gains as well as large losses were received when participants failed to respond within the allowable time window, so these two outcomes served as de facto negative RPEs.

| Criteria | Needs (hunger) | ‘Wanting’ |
| --- | --- | --- |
| Privation contrast | Yes | N/A |
| Presence of cue indicating the reward | Yes | Yes |
| Cue that triggers decision (to get the reward to be gained during the task) | Not necessarily | Yes |
| fMRI contrast taken only during the anticipation (cue) or after (during) the viewing of the cue | Yes | Yes |
| The reward is relevant for the need (self-specific) | Yes | N/A |
| Do not contrast two rewards | N/A (ex: High calorie - low calorie experiments were included) | Yes (with one exception) |
| Healthy individuals only | Yes (however, we took people that had up to 30-35 of BMI, especially in contrasts when healthy and overweight were mixed) | Yes |
| MNI or Talairach Coordinates | Yes | Yes |
| Whole brain contrast (with or without SVC) | Yes | Yes |
| Corrected | Yes | Yes |
| Activation contrast only | Yes | Yes |
| Excluded: MRI and resting states; cognitive conjunction analysis; and functional connectivity results | Yes | Yes |

**Table 2.** ‘Wanting’ST selected articles List of articles that were selected. for ‘Wanting’_ST_..

| Paper | Stimulus and Cue | Task description | Contrast | Healthy Participants |
| --- | --- | --- | --- | --- |
| Schneider, M., Leuchs, L., Czisch, M., Sämann, P. G., & Spormaker, V. I. (2018). Disentangling reward anticipation with simultaneous pupillometry/fMRI. *NeuroImage*, 178, 11-22. | Money, reward | MIDT and Pupillometry | Reward anticipation-control | 46 |
| Wu, C. C., Samanez-Larkin, G. R., Katovich, K., & Knutson, B. (2014). Affective traits link to reliable neural markers of incentive anticipation. *Neuroimage*, 84, 279-289. | Money | MIDT | Anticipation: Gain-Non Gain | 52 |
| Ubl, B., Kuehner, C., Kirsch, P., Rutter, M., Diener, C., & Flor, H. (2015). Altered neural reward and loss processing and prediction error signalling in depression. *Social cognitive and affective neuroscience*, 10(8), 1102-1112. | Money | monetary reward paradigm | Anticipation: high gain vs control | 28 |
| Jia, T., Macare, C., Desrivieres, S., Gonzalez, D. A., Tao, C., Ji, X., ... & Bokde, A. L. (2016). Neural basis of reward anticipation and its genetic determinants. *Proceedings of the National Academy of Sciences*, 113(14), 3879-3884. | Money | MIDT | Anticipation: high win vs. no win | 1544 |
| Young, C. B., & Nusslock, R. (2016). Positive mood enhances reward-related neural activity. *Social cognitive and affective neuroscience*, 11(6), 934-944. | Money | MIDT | Reward vs. Non-reward (anticipation) | 40 |
| Millman, Z. B., Gallagher, K., Demro, C., Schiffman, J., Reeves, G. M., Gold, J. M., ... & Buchanan, R. W. (2019). Evidence of reward system dysfunction in youth at clinical high-risk for psychosis from two event-related fMRI paradigms. *Schizophrenia research*. | Money | MIDT | Large Received Gain > Small Received Gain | 41 |
| Navas, J. F., Barrós-Loscertales, A., Costumero-Ramos, V., Verdejo-Román, J., Vilar-López, R., & Verdejo-García, A. (2018). Excessive body fat linked to blunted somatosensory cortex response to general reward in adolescents. *International Journal of Obesity*, 42(1), 88. | Money | MIDT | reward anticipation | 68 |
| Herbort, M. C., Soch, J., Wüstenberg, T., Krauel, K., Pujara, M., Koenigs, M., ... & Schott, B. H. (2016). A negative relationship between ventral striatal loss anticipation response and impulsivity in borderline personality disorder. *NeuroImage: Clinical*, 12, 724-736. | Money | MIDT | gain anticipation | 23 |
| Kohls, G., Perino, M. T., Taylor, J. M., Madva, E. N., Cayless, S. J., Troiani, V., ... & Schultz, R. T. (2013). The nucleus accumbens is involved in both the pursuit of social reward and the avoidance of social punishment. *Neuropsychologia*, 51(11), 2062-2069. | Social incentive | SIDT | Anticipation of social approval | 22 |
| Kumar, P., Berghorst, L. H., Nickerson, L. D., Dutra, S. J., Goer, F. K., Greve, D. N., & Pizzagalli, D. A. (2014). Differential effects of acute stress on anticipatory and consummatory phases of reward processing. *Neuroscience*, 266, 1-12. | Money | MIDT | Anticipation (Reward vs. No-incentive Cue) | 18 |
| Bradley, K. A., Case, J. A., Freed, R. D., Stern, E. R., & Gabbay, V. (2017). Neural correlates of RDoC reward constructs in adolescents with diverse psychiatric symptoms: A Reward Flanker Task pilot study. *Journal of affective disorders*, 216, 36-45. | Money | Reward Flanker Task | Reward Anticipation vs. Implicit Baseline | 22 |
| Richter, A., Petrovic, A., Diekhof, E. K., Trost, S., Wolter, S., & Gruber, O. (2015). Hyperresponsivity and impaired prefrontal control of the mesolimbic reward system in schizophrenia. *Journal of psychiatric research*, 71, 8-15. | Points, targets and CS | Desire-reason paradigm | desire context reason context | 16 |
| Gluth, S., Rieskamp, J., & Büchel, C. (2013). Neural evidence for adaptive strategy selection in value-based decision-making. *Cerebral Cortex*, 24(8), 2009-2021. | Investment | dynamic learning task | Expected value | 24 |
| Trost, S., Diekhof, E. K., Mohr, H., Vieker, H., Krämer, B., Wolf, C., ... & Gruber, O. (2016). Investigating the impact of a genome-wide supported bipolar risk variant of MAD1L1 on the human reward system. *Neuropsychopharmacology*, 41(11), 2679. | Points, targets and CS | Desire-reason paradigm | desire context reason context | 224 |
| Trost, S., Diekhof, E. K., Zvonik, K., Lewandowski, M., Usher, J., Keil, M., ... & Gruber, O. (2014). Disturbed anterior prefrontal control of the mesolimbic reward system and increased impulsivity in bipolar disorder. *Neuropsychopharmacology*, 39(8), 1914. | Points, targets and CS | Desire-reason paradigm | desire context | 16 |
| Yu, R., Mobbs, D., Seymour, B., Rowe, J. B., & Calder, A. J. (2014). The neural signature of escalating frustration in humans. *Cortex*, 54, 165-178. ISO 690 | cue, and coin | multi-trial reward schedule task | Cue – block Cue (increased proximity) Cue (increased expended effort) | 27 |
| Krebs, R. M., Schott, B. H., Schütze, H., & Düzel, E. (2009). The novelty exploration bonus and its attentional modulation. *Neuropsychologia*, 47(11), 2272-2281. | cue, reward | number comparison task (NCT) | reward-predicting cues in Exp 1: Contrast reward vs. neutral familiar reward-predicting cues exp 1 reward-predicting cues in Exp 2: Contrast reward vs. neutral novel reward-predicting cues exp 2 | 24 exp1<br>20 exp 2 |
| Articles from other sources |  |  |  |  |
| Samanez-Larkin, G. R., Gibbs, S. E., Khanna, K., Nielsen, L., Carstensen, L. L., & Knutson, B. (2007). Anticipation of monetary gain but not loss in healthy older adults. *Nature neuroscience*, 10(6), 787. | Money | MIDT | Gain vs nongain anticipation: younger<br>Gain vs nongain anticipation: older | 12<br>12 |
| Simon, J. J., Walther, S., Fiebach, C. J., Friederich, H. C., Stippich, C., Weisbrod, M., & Kaiser, S. (2010). Neural reward processing is modulated by approach-and avoidance-related personality traits. *Neuroimage*, 49(2), 1868-1874. | Money | MIDT | Anticipation of reward vs. non-reward | 24 |
| Wittmann, B. C., Schott, B. H., Guderian, S., Frey, J. U., Heinze, H. J., & Düzel, E. (2005). Reward-related FMRI activation of dopaminergic midbrain is associated with enhanced hippocampus-dependent long-term memory formation. *Neuron*, 45(3), 459-467. | Money | MIDT | Reward Anticipation | 16 |
| Murray, L., Lopez-Duran, N. L., Mitchell, C., Monk, C. S., & Hyde, L. W. (2020). Neural mechanisms of reward and loss processing in a low-income sample of at-risk adolescents. *Social cognitive and affective neuroscience*, 15(12), 1299-1314. | Points | Lottery choice task | reward anticipation> neutral anticipation | 128 |
| Yao, Y. W., Liu, L., Worhunsky, P. D., Lichenstein, S., Ma, S. S., Zhu, L., ... & Yip, S. W. (2020). Is monetary reward processing altered in drug-naïve youth with a behavioral addiction? Findings from internet gaming disorder. *NeuroImage: Clinical*, 26, 102202. | Money | MID task | Gain anticipation | 27 |

Regarding ‘Needing’_ST_ related articles, we typed the keywords on PubMed (February 2021). The database returned 376 articles. The main logic was to select experiments when subjects were in a hungry state and perceiving a food stimulus. We looked for both “hunger>baseline” as well as “hunger>satiety” contrasts, because of the inherent subtraction logic of fMRI and in order to have a larger number of experiments. Hence, the two main criteria were : 1) presence of a privation contrast: hunger +stimulus > satiety +stimulus, or hunger+stimulus > baseline; 2) the participant was perceiving some food stimulus which could be presented in any modality : visual, taste, odor, etc. Using the selection criteria (see table 1 for all criteria) we kept 26 articles. After fully reading the final 26 articles, nine were selected, and we found some additional ones through other articles and reviews, and seven among them matched all criteria for ‘Needing’_ST_ (hunger) for a total of 16 articles (see table 3 for list of retained articles). See Prisma in supplementary material for step by step exclusion of articles.

**Table 3.** ‘Needing’ST selected articles List of articles that were selected. for ‘Needing’_ST_..

| Paper | Contrast | Stimuli | Task | Healthy Participants | Fasted hours |
| --- | --- | --- | --- | --- | --- |
| Jiang, T., Soussignan, R., Schaal, B., & Royet, J. P. (2014). Reward for food odors: an fMRI study of liking and wanting as a function of metabolic state and BMI. Social cognitive and affective neuroscience, 10(4), 561-568. | Liking – Wanting (hunger)<br>Wanting – Liking (hunger)<br>Food - NFood (hunger) | Odor | Odor presentation and rating | 12 |  |
| Green, E., Jacobson, A., Haase, L., & Murphy, C. (2015). Neural correlates of taste and pleasantness evaluation in the metabolic syndrome. Brain research, 1620, 57-71. | Hunger-Satiety: Control<br>Sucrose>Caffeine (during hunger)<br>Caffeine>Sucrose (hunger) | Taste | Swallowing aqueous solution | 15 | 12h |
| Martens, M. J., Born, J. M., Lemmens, S. G., Karhunen, L., Heinecke, A., Goebel, R., ... & Westerterp-Plantenga, M. S. (2013). Increased sensitivity to food cues in the fasted state and decreased inhibitory control in the satiated state in the overweight. The American journal of clinical nutrition, 97(3), 471-479. | Fasted: F>NF<br>Fasted: stimuli - subject group<br>Fasted: correlation F>NF with BMI | Visual | Viewing food and non food pictures | 40 | 10h |
| LaBar, K. S., Gitelman, D. R., Parrish, T. B., Kim, Y. H., Nobre, A. C., & Mesulam, M. (2001). Hunger selectively modulates corticolimbic activation to food stimuli in humans. Behavioral neuroscience, 115(2), 493. | Hungry-Satiated | Visual | Viewing food and tool images | 17 | 8h |
| Harding, I. H., Andrews, Z. B., Mata, F., Orlandea, S., Martinez-Zalacain, I., Soriano-Mas, C., ... & Verdejo-Garcia, A. (2018). Brain substrates of unhealthy versus healthy food choices: influence of homeostatic status and body mass index. International Journal of Obesity, 42(3), 448. | Healthy versus unhealthy food choice: fasted>satiated | Visual | participants were asked to select an option using a two-button response box. | 30 | 10h |
| Frank, S., Laharnar, N., Kullmann, S., Veit, R., Canova, C., Hegner, Y. L., ... & Preissl, H. (2010). Processing of food pictures: influence of hunger, gender and calorie content. Brain research, 1350, 159-166. | HiCal hungry vs. HiCal satiated | Visual | One-back task: press a button, either to indicate that the seen image was the same or another button to indicate that the picture was not the same | 12 | 8h |
| Holsen, L. M., Zarcone, J. R., Thompson, T. I., Brooks, W. M., Anderson, | Premeal : Food > non-food | Visual | Viewing pictures of food, animals, and baseline | 9 | 4h |
| M. F., Ahluwalia, J. S., ... & Savage, C. R. (2005).<br>Neural mechanisms underlying food motivation in children and adolescents. *Neuroimage*, 27(3), 669-676. |  |  | control images |  |  |
| Jacobson, A., Green, E., Haase, L., Szajer, J., & Murphy, C. (2019).<br>Differential Effects of BMI on Brain Response to Odor in Olfactory, Reward and Memory Regions: Evidence from fMRI. *Nutrients*, 11(4), 926. | Odor during the hunger condition | Odor/Taste | Odor stimuli delivered to the tongue | 40 | 12h |
| Articles from other sources |  |  |  |  |  |
| Haase, L., Green, E., & Murphy, C. (2011). Males and females show differential brain activation to taste when hungry and sated in gustatory and reward areas. *Appetite*, 57(2), 421-434. | Hunger x male x Sucrose > water<br>Hunger x female x NaCl > water<br>Hunger x female x CAFFEINE > water<br>Hunger x female x SUCROSE > water<br>Hunger x female x CITRIC ACID > water | Taste | Taste stimuli presentation | 21 | 12h |
| Führer, D., Zysset, S., & Stumvoll, M. (2008). Brain | Hunger > Satiety | Visual | Viewing pictures and two-back | 12 | 14h |
| activity in hunger and satiety: an exploratory visually stimulated FMRI study. Obesity, 16(5), 945-950. |  |  | task: the subject was asked to press a button when the same letter was shown two steps earlier |  |  |
| Haase, L., Cerf-Ducastel, B., & Murphy, C. (2009). Cortical activation in response to pure taste stimuli during the physiological states of hunger and satiety. Neuroimage, 44(3), 1008-1021. | Hunger > satiety x sucrose<br>Hunger > satiety x Caffeine<br>Hunger > satiety x Saccharin<br>Hunger > satiety x Citric acid | Taste | Stimulus presentation delivered to the tip of the tongue | 18 | 12h |
| Uher, R., Treasure, J., Heining, M., Brammer, M. J., & Campbell, I. C. (2006). Cerebral processing of food-related stimuli: effects of fasting and gender. Behavioural brain research, 169(1), 111-119. | Fasting > satiety x food-related stimuli (Chocolate AND Chicken)<br>Fasting > satiety x food-related stimuli (Chocolate)<br>Fasting > satiety x food-related stimuli (Chicken) | Visual | Viewing photographs of food | 18 | 24h |
| Holsen, L. M., Zarcone, J. R., Brooks, W. M., Butler, M. G., Thompson, T. I., Ahluwalia, J. S., ... & Savage, C. R. (2006). Neural mechanisms underlying hyperphagia in Prader-Willi syndrome. | HW x Pre-meal x food > non-food<br>Hw x Pre-meal x non-food > food | Visual | Viewing pictures of food, animals and control | 9 | 4h |
| Obesity, 14(6), 1028-1037. |  |  |  |  |  |
| Jacobson, A., Green, E., & Murphy, C. (2010). Age-related functional changes in gustatory and reward processing regions: An fMRI study. *Neuroimage*, 53(2), 602-610. | Hunger x Older adults x Sucrose<br>Hunger x Young adults x Sucrose<br>Hunger x Older adults x Citric acid<br>Hunger x Younger adults x Citric acid<br>Hunger x Older adults x NaCl<br>Hunger x Younger adults x NaCl<br>Hunger x Older adults x Caffeine<br>Hunger x Younger adults x Caffeine | Taste | Stimuli were delivered orally | 38 | 12h |
| Cheah, Y. S., Lee, S., Ashoor, G., Nathan, Y., Reed, L. J., Zelaya, F. O., ... & Amiel, S. A. (2014). Ageing diminishes the modulation of human brain responses to visual food cues by meal ingestion. *International Journal of Obesity*, 38(9), 1186. | FASTED > FED x Visual food cue | Visual | Viewing food and non food | 24 | 8h |
| He, Q., Huang, X., Zhang, S., Turel, O., Ma, L., & Bechara, A. (2019). Dynamic causal modeling | Hungry > Satiated | Visual | Food pictures | 45 | 14-15h |
| of insular, striatal, and prefrontal cortex activities during a food-specific Go/NoGo task. *Biological Psychiatry: Cognitive Neuroscience and Neuroimaging*, 4(12), 1080-1089. |  |  |  |  |  |

### Meta-analyses

Meta-analyses were conducted with the activation likelihood estimation (ALE) approach using the Brainmap’s GingerALE application. Independently introduced by Turkeltaub and his colleagues (2002) and by Chein and colleagues (2002) and revised by Eickhoff and colleagues (2009), the ALE meta-analysis treats activation foci not as single point, but as spatial probability distributions that are centered at the given coordinates (Eickhoff et. al, 2012). The Eickhoff and colleagues’s revised ALE algorithm (2009) models the spatial uncertainty by using an estimation of the inter-subject and inter-laboratory variability (which is typically observed in neuroimaging experiments). Then, union of activation probabilities for each voxel of all included experiment is computed to give an ALE map; and a permutation procedure (in which data sets are created similar to the real data in terms of number of experiments, foci per experiments and number of subjects, but in which foci are randomly distributed) is used in order to test the differentiation between true convergence of foci and random clustering (Eickhoff, et. al, 2012). As a method of inference, the new algorithm uses random-effects analysis that calculates the above-chance clustering between experiments. Furthermore, the new algorithm gives more weight to grey matter compared to white matter by limiting the meta-analysis to an anatomically constrained space specified by a grey matter mask. Contrasts analyses are based on two different datasets (i.e. two previous ALE results) and thus compare two different sets of foci for statistically significant differences; and the conjunction is the intersection of the thresholded maps.

In our analyses we used the MNI152 coordinate system and the less conservative (larger) mask size. For ‘Wanting’_ST_, there were 21 articles, 34 experiments, 3306 subjects and 572 foci. (See table 2 and 3 for all included articles.). For ‘Needing’_ST,_ (hunger), we had 16 articles, 38 experiments, 733 subjects and 494 foci. In our study, for main individual meta-analyses, all maps were thresholded using a cluster-level family-wise error (cFWE) correction (P < 0.05) with a cluster-forming threshold of P < 0.001(uncorrected at the voxel level) (Eklund et al., 2016; Woo et al., 2014), and 1000 permutations. For the contrast meta-analyses we used the two cFWE corrected maps with p < 0.01 (uncorrected at the voxel level), 10,000 permutations (see Eickhoff et. al., 2011); and the conjunction was the intersection of the two cFWE thresholded maps. Maps from meta-analyses were overlaid on a MNI template and viewed using Mango (http://ric.uthscsa.edu/mango/).

Our inclusion criteria (such as including only corrected results and experiments) lowered the number of included experiments. Thus, to confirm that our main meta-analytic results were not driven by the coordinates from a single publication, we conducted validation analyses using a leave-one-experiment-out (LOEO) approach. In this approach, on each fold, one contrast (i.e., experiment) was excluded and the ALE meta-analysis was conducted on the remaining N – 1 contrasts. Thus, results from this procedure consisted of brain regions that were identified in every fold of the LOEO, and are not mainly driven by a single contrast.

## RESULTS

### Main meta-analyses

#### ‘‘Wanting’_ST_

Our first meta-analysis was on ‘Wanting’_ST_ *(table 4 and figure 1)*. This meta-analysis revealed consistent activations within the following regions: the left putamen, the left globus pallidus (which encompassed the nucleus accumbens), the left caudate body and right caudate head, the left substantia nigra, the right red nucleus (encompassing the ventral tegmental area), the right hypothalamus, the bilateral thalamus, the left precentral gyrus, the left inferior parietal lobule, the right dorso-lateral and medial prefrontal cortex, the right superior parietal lobule, the right claustrum ( whose cluster was mainly the anterior insula).

**Table 4.** ‘Wanting’_ST_ Coordinates for peak activated clusters in the ‘Wanting’_ST_ condition.

| Cluster # | x | y | z | ALE | P | Z | Label (Nearest Gray Matter within 5mm) |
| --- | --- | --- | --- | --- | --- | --- | --- |
| 1 | -16 | 8 | 2 | 0.073149 | 8.39E-21 | 9.281936 | Left Cerebrum.Sub-lobar.Lentiform Nucleus.Gray Matter.Putamen |
| 1 | -32 | 18 | 4 | 0.072856 | 1.07E-20 | 9.256376 | Left Cerebrum.Sub-lobar.Clastrum.Gray Matter.* |
| 1 | 12 | 8 | 0 | 0.062949 | 3.04E-17 | 8.364339 | Right Cerebrum.Sub-lobar.Caudate.Gray Matter.Caudate Head |
| 1 | -18 | 0 | 14 | 0.041655 | 1.70E-10 | 6.279265 | Left Cerebrum.Sub-lobar.Caudate.Gray Matter.Caudate Body |
| 1 | 18 | -6 | 8 | 0.023581 | 1.41E-05 | 4.186929 | Right Cerebrum.Sub-lobar.Thalamus.Gray Matter.Ventral Anterior Nucleus |
| 1 | -10 | -6 | -2 | 0.022746 | 2.26E-05 | 4.078712 | Left Cerebrum.Sub-lobar.Thalamus.Gray Matter.* |
| 1 | 10 | -6 | -6 | 0.022589 | 2.48E-05 | 4.057735 | Right Cerebrum.Sub-lobar.*.Gray Matter.Hypothalamus |
| 1 | -16 | -4 | -2 | 0.019475 | 1.37E-04 | 3.639444 | Left Cerebrum.Sub-lobar.Lentiform Nucleus.Gray Matter.Medial Globus Pallidus |
| 1 | 8 | -6 | 10 | 0.018669 | 2.11E-04 | 3.525875 | Right Cerebrum.Sub-lobar.Thalamus.Gray Matter.Anterior Nucleus |
| 2 | -6 | -22 | -18 | 0.070495 | 7.37E-20 | 9.047366 | Left Brainstem.Midbrain.*.Gray Matter.Substantia Nigra |
| 2 | 6 | -22 | -18 | 0.062512 | 4.28E-17 | 8.323777 | Right Brainstem.Midbrain.*.Gray Matter.Red Nucleus |
| 3 | -6 | 8 | 52 | 0.055675 | 7.87E-15 | 7.681671 | Left Cerebrum.Frontal Lobe.Medial Frontal Gyrus.Gray Matter.Brodmann area 6 |
| 4 | 34 | 22 | 2 | 0.071349 | 3.69E-20 | 9.122826 | Right Cerebrum.Sub-lobar.Clastrum.Gray Matter.* |
| 5 | -28 | -8 | 56 | 0.036814 | 4.27E-09 | 5.757467 | Left Cerebrum.Frontal Lobe.Precentral Gyrus.Gray Matter.Brodmann area 6 |
| 6 | -44 | -36 | 46 | 0.02985 | 3.47E-07 | 4.963281 | Left Cerebrum.Parietal Lobe.Inferior Parietal Lobule.Gray Matter.Brodmann area 40 |
| 7 | 38 | 36 | 28 | 0.045997 | 8.58E-12 | 6.728507 | Right Cerebrum.Frontal Lobe.Middle Frontal Gyrus.Gray Matter.Brodmann area 9 |
| 8 | 28 | -4 | 50 | 0.03858 | 1.34E-09 | 5.950417 | Right Cerebrum.Frontal Lobe.Middle Frontal Gyrus.Gray Matter.Brodmann area 6 |
| 9 | 34 | -52 | 44 | 0.0408 | 3.04E-10 | 6.188297 | Right Cerebrum.Parietal Lobe.Superior Parietal Lobule.Gray Matter.Brodmann area 7 |
| 10 | -44 | 2 | 34 | 0.045089 | 1.62E-11 | 6.635556 | Left Cerebrum.Frontal Lobe.Precentral Gyrus.Gray Matter.Brodmann area 6 |
| 11 | -28 | 38 | 12 | 0.027964 | 1.09E-06 | 4.736878 | No Gray Matter found |
| 11 | -32 | 44 | 14 | 0.024484 | 8.43E-06 | 4.302969 | Left Cerebrum.Frontal Lobe.Middle Frontal Gyrus.Gray Matter.Brodmann area 10 |
| 11 | -36 | 50 | 22 | 0.019078 | 1.69E-04 | 3.583664 | Left Cerebrum.Frontal Lobe.Middle Frontal Gyrus.Gray Matter.Brodmann area 9 |
| 12 | 46 | -34 | 46 | 0.035456 | 1.02E-08 | 5.608079 | Right Cerebrum.Parietal Lobe.Inferior Parietal Lobule.Gray Matter.Brodmann area 40 |

**Fig. 1.**
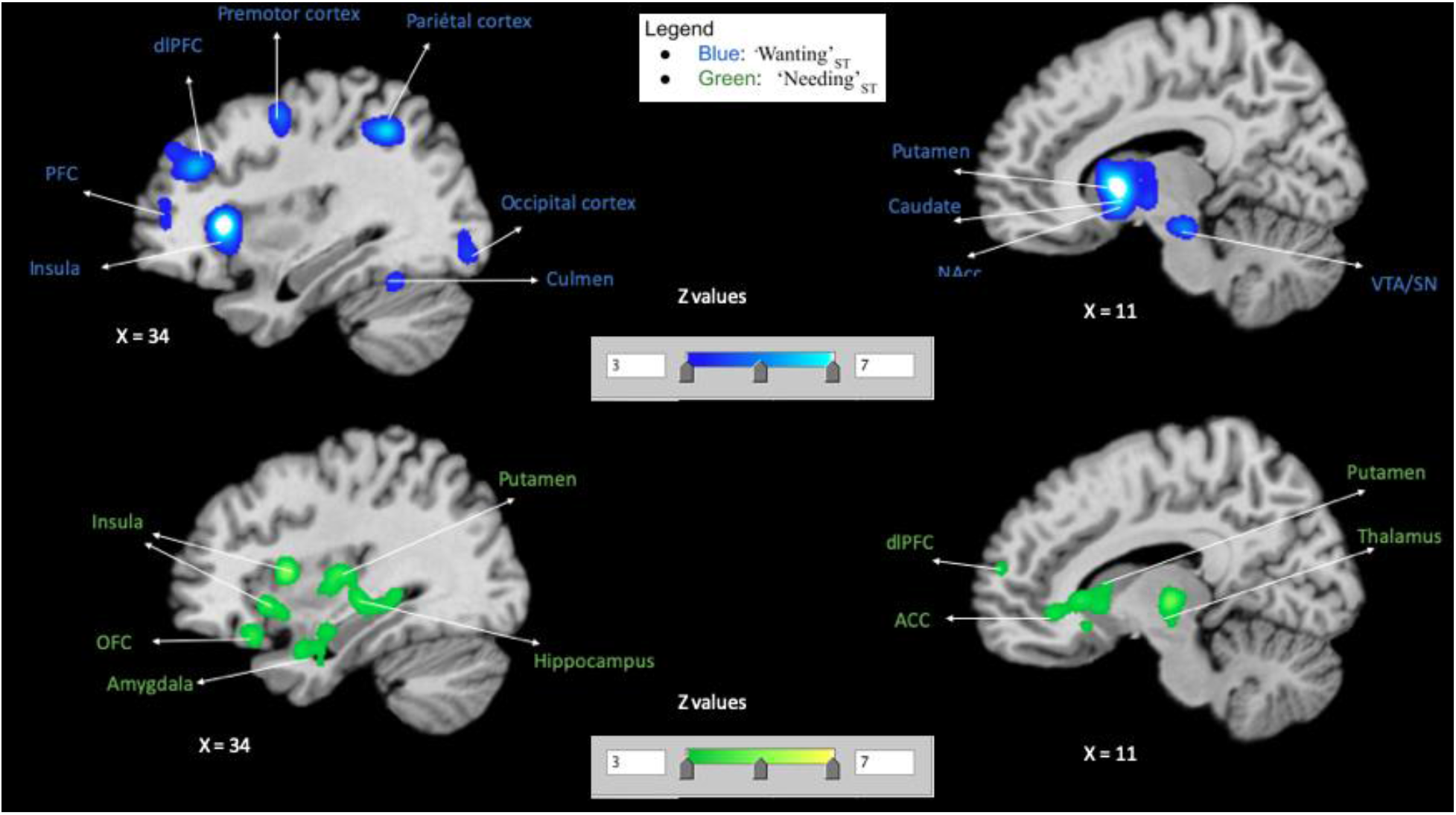
Single meta-analyses maps. Maps for activated clusters in each condition: ‘Wanting’_ST_ (blue) and ‘Needing’_ST_ (green), showing activation pattern for each.

#### ‘Needing’_ST_

Next, we conducted an individual meta-analysis on Needing’_ST_ (hunger with stimulus; *Table 5 and figure 1)*. This second meta-analysis revealed consistent activations in : the bilateral anterior insula, right middle and posterior insula, right thalamus, left claustrum, right hippocampus, bilateral putamen, right caudate body, right caudate head (encompassing the NAcc), and right posterior putamen (encompassing the caudate tail), amygdala, bilateral anterior cingulate area (encompassing the right OFC), right uncus and left subcallosal area (which can be considered as entorhinal cortex (Fischl et al., 2009), and the right mammillary body.

**Table 5.** ‘Needing’_ST_ Coordinates for peak activated clusters in the ‘Needing’_ST_ condition.

| Cluster # | x | y | z | ALE | P | Z | Label (Nearest Gray Matter within 5mm) |
| --- | --- | --- | --- | --- | --- | --- | --- |
| 1 | 38 | 10 | 6 | 0.030085 | 9.63E-09 | 5.61851 | Right Cerebrum.Sub-lobar.Insula.Gray Matter.Brodmann area 13 |
| 1 | 14 | -16 | 0 | 0.027834 | 5.17E-08 | 5.32083 | Right Cerebrum.Sub-lobar.Thalamus.Gray Matter.Mammillary Body |
| 1 | 42 | -4 | -2 | 0.022411 | 2.42E-06 | 4.571283 | Right Cerebrum.Sub-lobar.Clastrum.Gray Matter.* |
| 1 | 44 | 2 | -10 | 0.022346 | 2.53E-06 | 4.56251 | Right Cerebrum.Sub-lobar.Insula.Gray Matter.Brodmann area 13 |
| 1 | 48 | -12 | 4 | 0.021699 | 3.97E-06 | 4.467 | Right Cerebrum.Sub-lobar.Insula.Gray Matter.Brodmann area 13 |
| 1 | 32 | -38 | -4 | 0.021315 | 5.15E-06 | 4.410615 | Right Cerebrum.Temporal Lobe.Sub-Gyral.Gray Matter.Hippocampus |
| 1 | 34 | -12 | 6 | 0.020947 | 6.64E-06 | 4.355373 | Right Cerebrum.Sub-lobar.Lentiform Nucleus.Gray Matter.Putamen |
| 1 | 34 | 18 | -8 | 0.020895 | 6.92E-06 | 4.346396 | Right Cerebrum.Sub-lobar.Clastrum.Gray Matter.* |
| 1 | 36 | -22 | -4 | 0.020609 | 8.37E-06 | 4.304496 | Right Cerebrum.Sub-lobar.Lentiform Nucleus.Gray Matter.Putamen |
| 1 | 28 | -8 | 0 | 0.019476 | 1.78E-05 | 4.134444 | Right Cerebrum.Sub-lobar.Lentiform Nucleus.Gray Matter.Putamen |
| 1 | 30 | -50 | -12 | 0.016798 | 9.81E-05 | 3.723894 | Right Cerebellum.Anterior Lobe.Culmen.Gray Matter.* |
| 2 | 14 | 36 | -4 | 0.022022 | 3.18E-06 | 4.514158 | Right Cerebrum.Limbic Lobe.Anterior Cingulate.Gray Matter.Brodmann area 24 |
| 2 | 10 | 24 | 0 | 0.021439 | 4.75E-06 | 4.428455 | Right Cerebrum.Sub-lobar.Caudate.Gray Matter.Caudate Head |
| 2 | 10 | 14 | -2 | 0.017611 | 5.89E-05 | 3.850592 | Right Cerebrum.Sub-lobar.Caudate.Gray Matter.Caudate Head |
| 2 | 8 | 20 | -10 | 0.017451 | 6.52E-05 | 3.825874 | Right Cerebrum.Limbic Lobe.Anterior Cingulate.Gray Matter.Brodmann area 25 |
| 2 | 12 | 16 | 4 | 0.017117 | 8.02E-05 | 3.774332 | Right Cerebrum.Sub-lobar.Caudate.Gray Matter.Caudate Body |
| 2 | 14 | 12 | 8 | 0.016643 | 1.08E-04 | 3.698454 | Right Cerebrum.Sub-lobar.Caudate.Gray Matter.Caudate Body |
| 2 | 14 | 22 | -12 | 0.015113 | 2.87E-04 | 3.443822 | Right Cerebrum.Sub-lobar.Caudate.Gray Matter.Caudate Head |
| 3 | -38 | 0 | -2 | 0.030925 | 5.07E-09 | 5.728328 | Left Cerebrum.Sub-lobar.Clastrum.Gray Matter.* |
| 3 | -40 | 8 | 12 | 0.017386 | 6.77E-05 | 3.816551 | Left Cerebrum.Sub-lobar.Insula.Gray Matter.Brodmann area 13 |
| 4 | 26 | 6 | -16 | 0.021686 | 3.99E-06 | 4.465524 | Right Cerebrum.Sub-lobar.Lentiform Nucleus.Gray Matter.Putamen |
| 4 | 30 | -10 | -18 | 0.02053 | 8.83E-06 | 4.292494 | Right Cerebrum.Limbic Lobe.Parahippocampal Gyrus.Gray Matter.Amygdala |
| 4 | 32 | -4 | -26 | 0.019465 | 1.80E-05 | 4.131432 | Right Cerebrum.Limbic Lobe.Parahippocampal Gyrus.Gray Matter.Amygdala |
| 4 | 34 | 4 | -28 | 0.019396 | 1.87E-05 | 4.122383 | Right Cerebrum.Limbic Lobe.Uncus.Gray Matter.Brodmann area 28 |
| 5 | -14 | 18 | -20 | 0.020973 | 6.55E-06 | 4.358362 | Left Cerebrum.Frontal Lobe.Medial Frontal Gyrus.Gray Matter.Brodmann area 25 |
| 5 | -26 | 4 | -16 | 0.018197 | 4.06E-05 | 3.940716 | Left Cerebrum.Frontal Lobe.Subcallosal Gyrus.Gray Matter.Brodmann area 34 |
| 5 | -14 | 22 | -12 | 0.017613 | 5.89E-05 | 3.850592 | Left Cerebrum.Sub-lobar.Caudate.Gray Matter.Caudate Head |
| 5 | -18 | 10 | -18 | 0.016592 | 1.12E-04 | 3.690466 | Left Cerebrum.Sub-lobar.Lentiform Nucleus.Gray Matter.Putamen |
| 5 | -6 | 22 | -10 | 0.015431 | 2.34E-04 | 3.498593 | Left Cerebrum.Limbic Lobe.Anterior Cingulate.Gray Matter.Brodmann area 24 |
| 6 | -34 | -22 | -4 | 0.025242 | 3.35E-07 | 4.970016 | Left Cerebrum.Sub-lobar.Lentiform Nucleus.Gray Matter.Putamen |

### Validation Results (LOEO Analyses)

The key output from the LOEO analysis was related to the robustness per cluster. That is, in what probability percentage a given cluster was observed. Here, we show from the LOEO analysis brain regions that have 100% probability of being activated in all experiments included in the meta-analyses.

#### ‘‘Wanting’_ST_ (supplementary material and figure 2)

For ‘Wanting’_ST_, consistent activations were identified in ALE-LOEO with 100% probability in the following peak regions: right midbrain (VTA and SN), right putamen (that included the caudate and the NAcc), left ACC, left caudate, left OFC, left anterior insula, left Inferior Parietal Lobule.

**Fig. 2.**
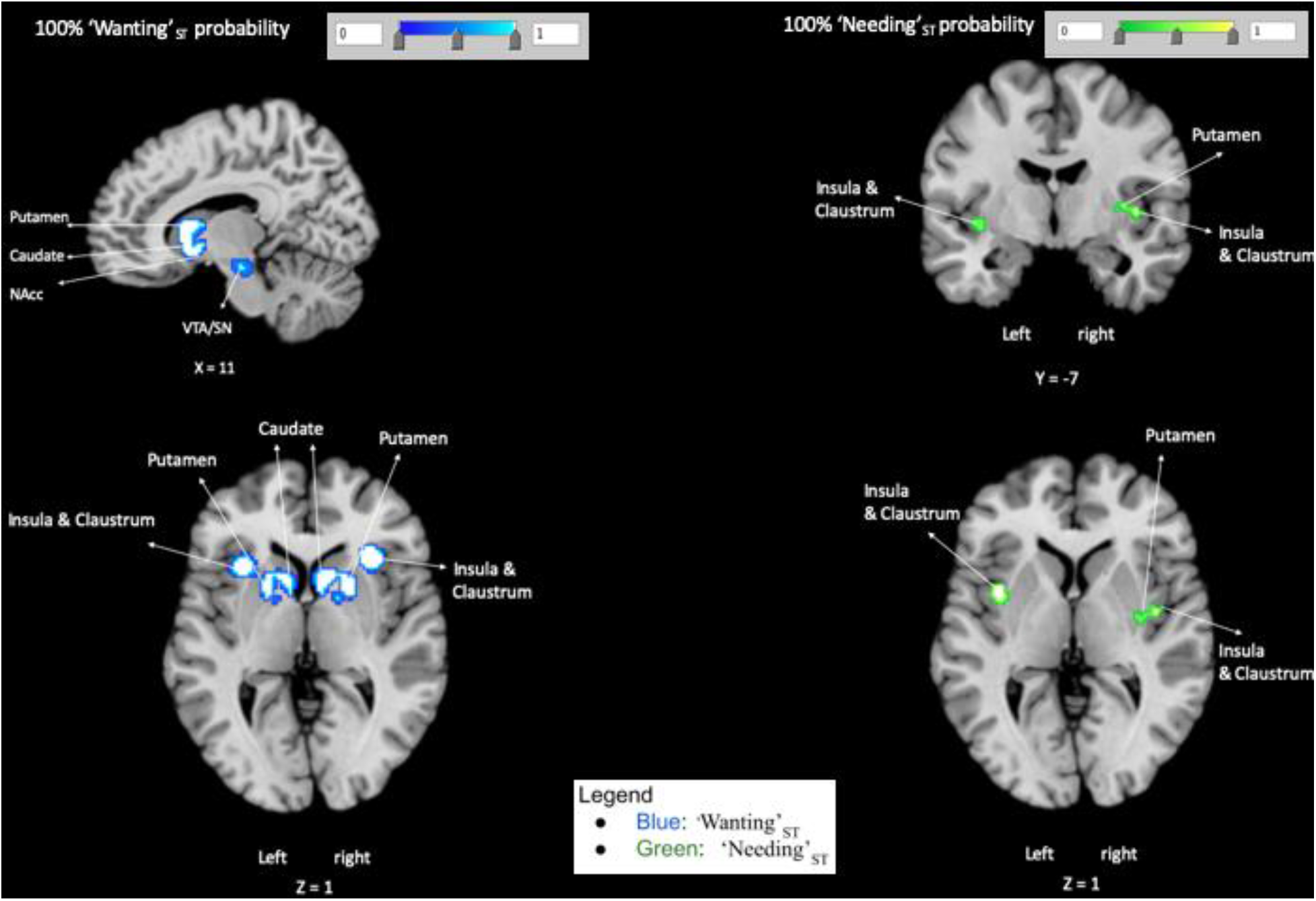
100% probability maps. LOEO Maps for clusters of ‘Wanting’_ST_ (blue) activation pattern and ‘Needing’_ST_ (green) activation pattern that have 100% probability of being activated in each included experiments.

#### ‘Needing’_ST_ (supplementary material and figure 2)

Regarding, ‘Needing’_ST_ ALE-LOEO meta-analysis revealed 3 peaks with 100% consistent activations in all experiments, within the bilateral middle insula, that included the caudoventral putamen and the claustrum.

### Conjunction and contrasts meta-analyses

Contrasts and conjunction analyses were based on ALE results of the two previous ALE results (‘Wanting’_ST_ AND ‘Needing’_ST_) that were compared for statistically significant differences and similarities.

#### ‘Wanting’_ST_ AND ‘Needing’_ST_ conjunction

The conjunction between ‘Wanting’_ST_ AND ‘Needing’_ST_ resulted in consistent activations within the head and body of the right caudate nucleus (the activated region does not include the nucleus accumbens), right claustrum and right anterior insula *(Table 6 and figure 3)*.

**Table 6.** ‘Wanting’_ST_ AND ‘Needing’_ST_ conjunction Coordinates for peak activated clusters in the ‘Wanting’_ST_ AND ‘Needing’_ST_ conjunction.

| Cluster # | x | y | z | ALE | Label (Nearest Gray Matter within 5mm) |
| --- | --- | --- | --- | --- | --- |
| 1 | 10 | 14 | -2 | 0.017611 | Right Cerebrum.Sub-lobar.Caudate.Gray Matter.Caudate Head |
| 1 | 12 | 16 | 4 | 0.017117 | Right Cerebrum.Sub-lobar.Caudate.Gray Matter.Caudate Body |
| 1 | 14 | 12 | 8 | 0.016643 | Right Cerebrum.Sub-lobar.Caudate.Gray Matter.Caudate Body |
| 2 | 34 | 20 | -6 | 0.020654 | Right Cerebrum.Sub-lobar.Clastrum.Gray Matter.* |
| 3 | 36 | 16 | 6 | 0.013647 | Right Cerebrum.Sub-lobar.Insula.Gray Matter.Brodmann area 13 |

**Fig. 3.**
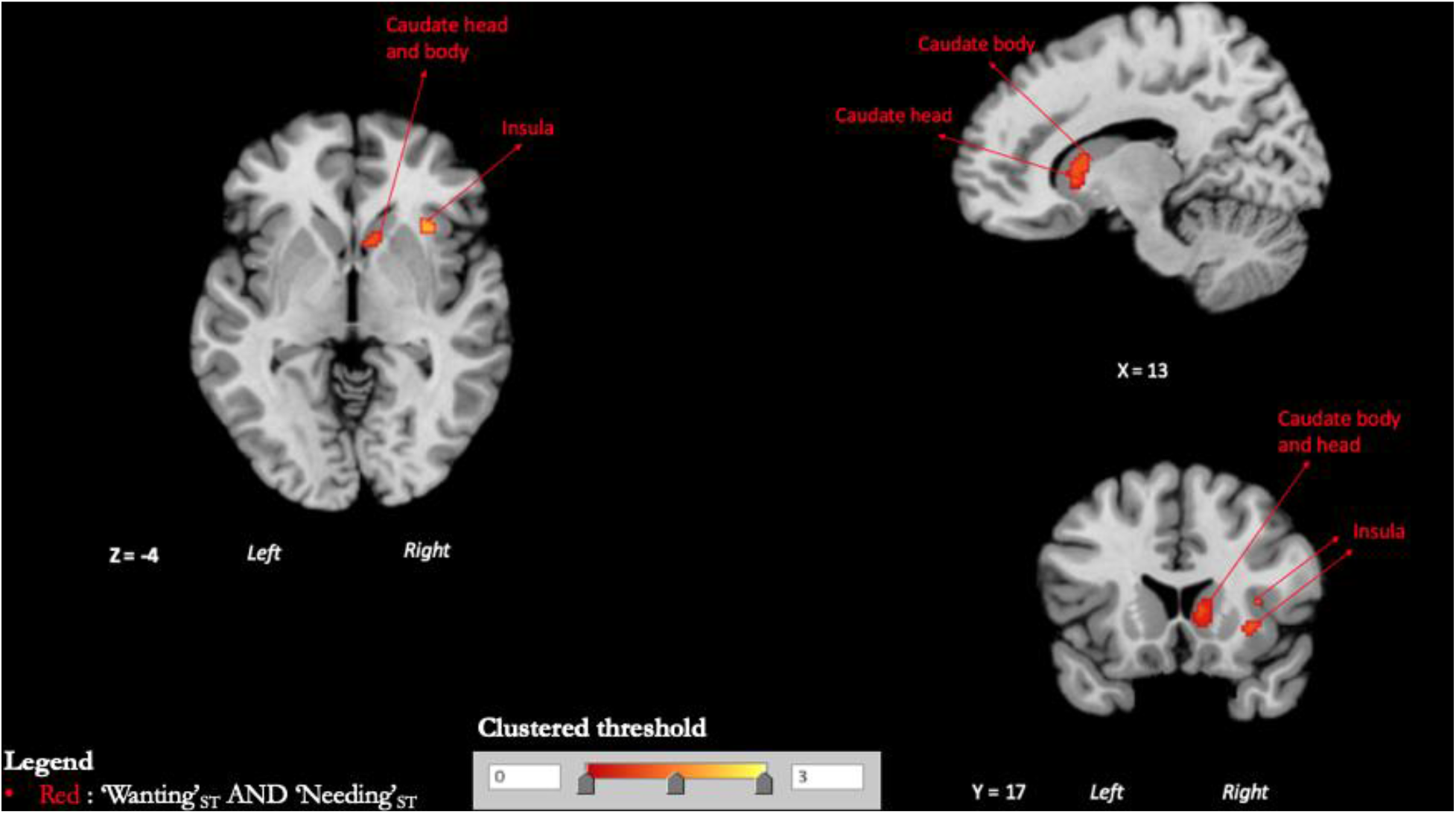
Conjunction maps. Clustered thresholded maps showing the intersection between activation patterns of‘Wanting’_ST_ and ‘Needing’_ST_.

#### Contrast: ‘Wanting’_ST_ -‘Needing’_ST_

Compared to ‘Needing’_ST_, *‘*Wanting’_ST_ more consistently activated regions of the left lateral globus pallidus (which encompassed the nucleus accumbens), the left red nucleus (encompassing the ventral tegmental area), right substantia nigra (SN), bilateral putamen, left anterior insula, the left precentral gyrus, the right superior parietal lobule, the left inferior parietal lobule, the right claustrum, the left anterior dorsolateral prefrontal cortex, and the right angular gyrus *(Table 7 and figure 4)*.

**Table 7.** Contrast: ‘Wanting’_ST_ -‘Needing’_ST_ Coordinates for peak activated clusters in the ‘Wanting’_ST_ =‘Needing’_ST_ contrast

| Cluster # | x | y | z | P | Z | Label (Nearest Gray Matter within 5mm) |
| --- | --- | --- | --- | --- | --- | --- |
| 1 | -16.4 | 9.4 | 3.2 | 0 | 3.890594 | Left Cerebrum.Sub-lobar.Lentiform Nucleus.Gray Matter.Putamen |
| 1 | -19.3 | 6 | 5.3 | 1.00E-04 | 3.719017 | Left Cerebrum.Sub-lobar.Lentiform Nucleus.Gray Matter.Putamen |
| 1 | -27.3 | 22 | 6.7 | 2.00E-04 | 3.540084 | Left Cerebrum.Sub-lobar.Clastrum.Gray Matter.* |
| 1 | -32.4 | 19.2 | 4.8 | 3.00E-04 | 3.431614 | Left Cerebrum.Sub-lobar.Insula.Gray Matter.Brodman area 13 |
| 2 | 15 | 5.5 | -2.5 | 0 | 3.890594 | Right Cerebrum.Sub-lobar.Lentiform Nucleus.Gray Matter.Lateral Globus Pallidus |
| 2 | 21 | 1 | 9 | 3.00E-04 | 3.431614 | Right Cerebrum.Sub-lobar.Lentiform Nucleus.Gray Matter.Putamen |
| 3 | -0.7 | -23 | -19.1 | 0 | 3.890594 | Left Brainstem.Midbrain.*.Gray Matter.Red Nucleus |
| 3 | 8 | -17 | -21 | 1.00E-04 | 3.719017 | No Gray Matter found |
| 3 | 11 | -21 | -20 | 2.00E-04 | 3.540084 | Right Brainstem.Midbrain.*.Gray Matter.Substantia Nigra |
| 4 | -2.6 | 7.9 | 52.2 | 0 | 3.890594 | Left Cerebrum.Frontal Lobe.Medial Frontal Gyrus.Gray Matter.Brodman area 6 |
| 5 | -30.6 | -8.8 | 59.1 | 0.00E+00 | 3.890594 | Left Cerebrum.Frontal Lobe.Precentral Gyrus.Gray Matter.Brodman area 6 |
| 6 | 29.7 | 21.6 | 2.2 | 0 | 3.890594 | Right Cerebrum.Sub-lobar.Clastrum.Gray Matter.* |
| 7 | -43.2 | -35.3 | 45 | 0 | 3.890594 | Left Cerebrum.Parietal Lobe.Inferior Parietal Lobule.Gray Matter.Brodman area 40 |
| 8 | 37 | 34.6 | 29.2 | 0 | 3.890594 | Right Cerebrum.Frontal Lobe.Middle Frontal Gyrus.Gray Matter.Brodman area 9 |
| 9 | 29.2 | -4.7 | 49.9 | 0 | 3.890594 | Right Cerebrum.Frontal Lobe.Middle Frontal Gyrus.Gray Matter.Brodman area 6 |
| 10 | 34.6 | -48.1 | 45 | 0 | 3.890594 | Right Cerebrum.Parietal Lobe.Superior Parietal Lobule.Gray Matter.Brodman area 7 |
| 10 | 34.2 | -56.4 | 44.2 | 1.00E-04 | 3.719017 | Right Cerebrum.Parietal Lobe.Angular Gyrus.Gray Matter.Brodman area 39 |
| 11 | -45.3 | 0.9 | 35.5 | 0 | 3.890594 | Left Cerebrum.Frontal Lobe.Precentral Gyrus.Gray Matter.Brodman area 6 |
| 12 | 44.8 | -37.5 | 44.8 | 0 | 3.890594 | Right Cerebrum.Parietal Lobe.Inferior Parietal Lobule.Gray Matter.Brodman area 40 |
| 12 | 46.3 | -29.7 | 45.4 | 1.00E-04 | 3.719017 | Right Cerebrum.Parietal Lobe.Inferior Parietal Lobule.Gray Matter.Brodmann area 40 |
| 13 | -28 | 44 | 14 | 0.0039 | 2.660607 | Left Cerebrum.Frontal Lobe.Middle Frontal Gyrus.Gray Matter.Brodmann area 10 |

**Fig. 4.**
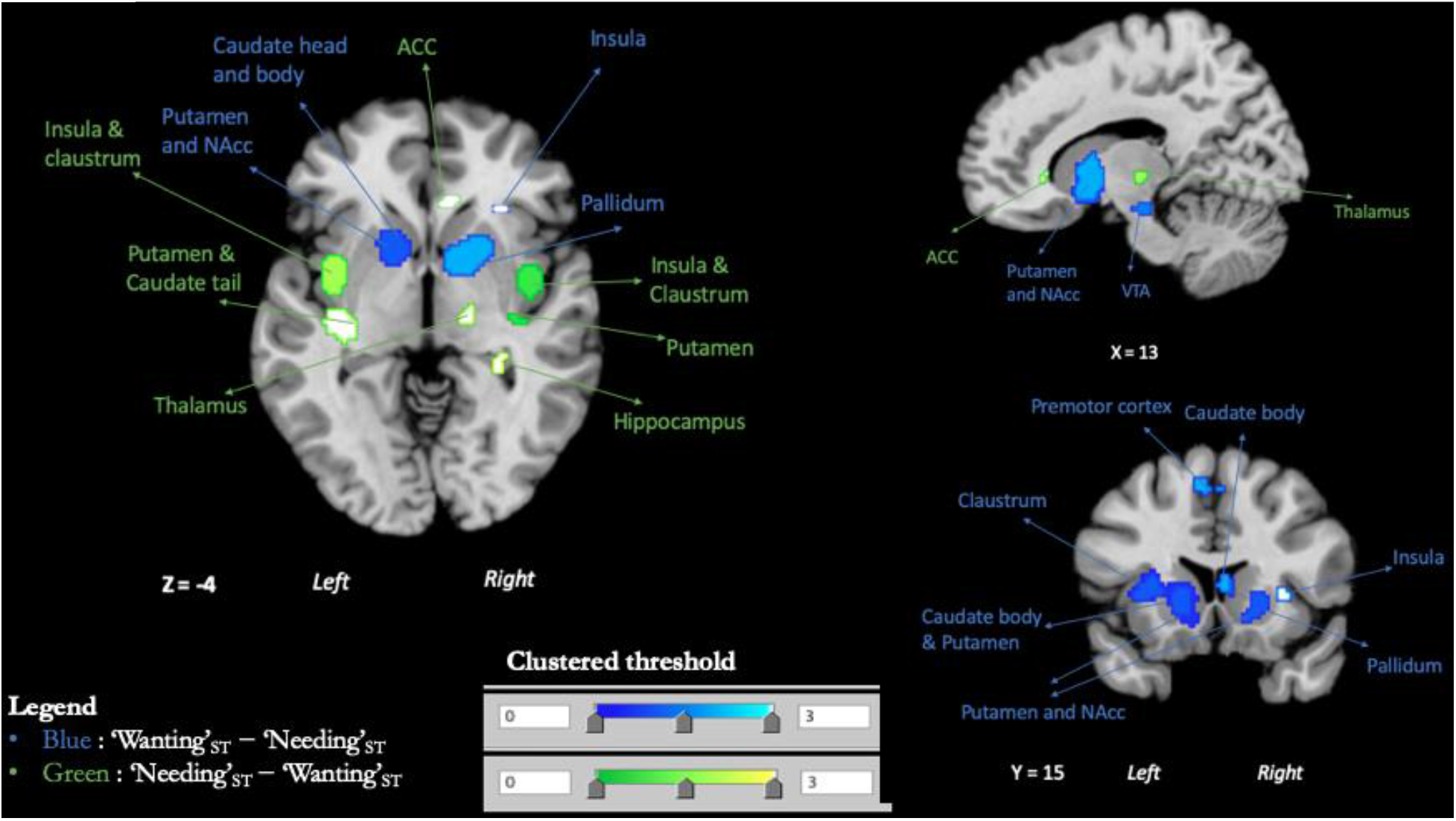
Contrasts maps. In blue, clustered thresholded maps for clusters of subtraction {[‘Wanting’_ST_] minus [‘Needing’_ST_]}. In green, clustered thresholded maps for clusters of subtraction {[‘Needing’_ST_] minus [‘Wanting’_ST_]}.

#### Contrast : ‘Needing’_ST_ -‘Wanting’_ST_

Compared to ‘Wanting’_ST_, ‘Needing’_ST_ more consistently activated regions of the right mid-posterior insula, bilateral claustrum, left putamen (encompassing the tail of caudate), right anterior cingulate area, right thalamus and bilateral hippocampus *(Table 8 and figure 4)*.

**Table 8.**
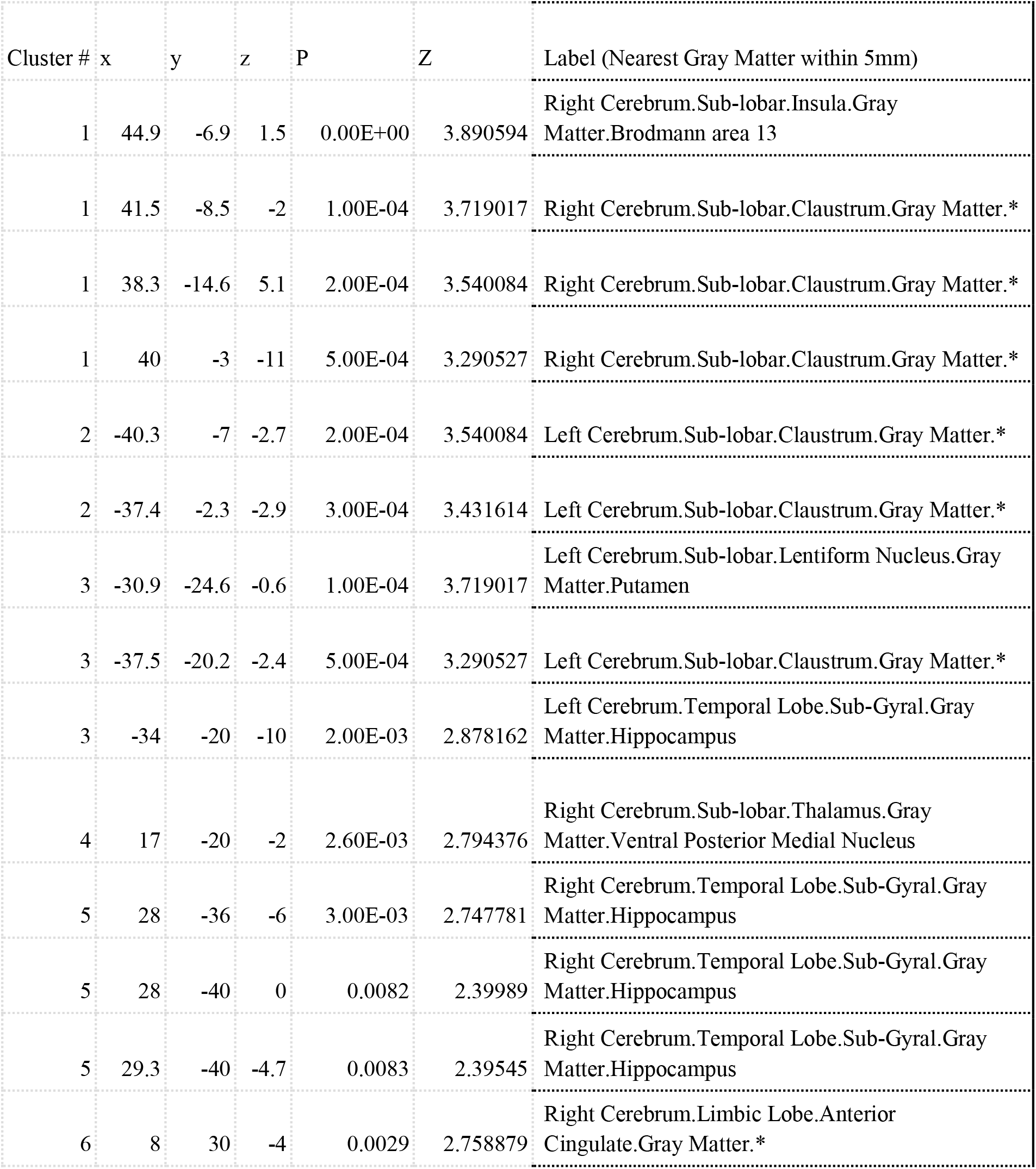
Contrast : ‘Needing’_ST_ -‘Wanting’_ST_ Coordinates for peak activated clusters in the ‘Needing’_ST_ -‘Wanting’_ST_ contrast

## DISCUSSION

Our goal was to compare the brain activation patterns related to value that comes from the state of ‘Wanting’_ST_ from the one from the state of ‘Needing’_ST_. To answer this, we used ALE neuroimaging meta-analysis, comparing consistent brain activation patterns during processing of stimuli in these two states. We used the perception of a cue predicting a reward for ‘Wanting’_ST_; and we used the perception of food stimuli in a hungry state as a model for ‘Needing’_ST_. We first carried out separate meta-analyses on Wanting’_ST_ and on ‘Needing’_ST_, then we contrasted and intersected them to identify differences and similarities between each of these states. We show that processing a stimulus in a ‘Wanting’_ST_ state seems more related to activity within the mesolimbic dopaminergic brain areas, nigrostriatal dopaminergic regions, and striatal regions; while processing a stimulus in a ‘Needing’_ST_ state seems more related to activity in viscerosensory cortices (i.e. mid-posterior insula) and caudal-ventral putamen (and to some extent the caudate tail). Both states seemed to share consistent activation in the caudate nucleus (head and body) and anterior insula. Compared to ‘Needing’_ST_, ‘Wanting’_ST_ more consistently activated the mesolimbic dopamine: the VTA and ventral striatum and pallidum, and nigrostriatal dopamine regions (i.e. SN and dorsal striatum). Compared to ‘Wanting’_ST_, ‘Needing’_ST_ more consistently activated the mid-posterior insula and ACC, caudo-ventral putamen and Caudate tail, and hippocampus. In the following paragraphs, we will discuss our overall results (the ones consistently found in all our meta-analyses) and how by identifying the brain areas most implicated for each state (‘Wanting’_ST_ vs ‘Needing’_ST_) can help us understand how we attribute different types of value to stimuli.

### Overview of consistent activation patterns for Wanting vs. Needing

Overall, our results--from main individual meta-analyses, LOEO analyses, and contrasts--confirm that the activation pattern of ‘Wanting’_ST_ related value shows consistent activation of VTA, ventral striatum, putamen, pallidum and anterior insula. Our results are in line with previous human studies using a wide range of methods or approaches (O’Doherty et. al, 2002; O’Doherty, 2004; Krebs et. al., 2009; Carter et, al., 2009; Schott et. al., 2008; Oldham et. al., 2018; Knutson et. al 2003; Knutson et. al 2001; Wilson et al., 2018; Simon et. al., 2015). For ‘Needing’_ST_, our results show that only the middle insula and to some extent the caudal-ventral putamen are consistently related to ‘Needing’_ST_ related value. The implication of the insula and dorsal striatum in ‘Needing’_ST_ is in accordance with some previous literature findings (Goldstone, et. al., 2009; van der Laan, et. al., 2011; Siep et. al., 2009). However, previous meta-analyses and studies on ‘Needing’_S_ had also identified other regions such as OFC, ACC and amygdala/parahippocampal gyrus (Führer et. al., 2008; LaBar et. al., 2011; Mohanty, et. al., 2008; Chen & Zeffiro, 2020). This could be due to the fact that we report here, regions that have been consistently found in all our meta-analyses (main, contrasts, and LOEO) and thus use a more stringent approach than in previous meta-analyses. Indeed, when only looking at results from our main meta-analysis, we also identified regions within the OFC, ACC, anterior insula and the amygdala. Nevertheless, using a more stringent approach, our results showing consistent activation mainly restricted to the mid-posterior insula make sense as it is often considered as the core viscerosensory cortex because it projects to other visceromotor cortices (anterior insula, OFC, ACC) (Barrett & Simmon, 2015); and dense multimodal sensory interoceptive prediction errors converge within the posterior insula to guide interoception (Gerlach et al., 2020). Thus, by combining contrasts, individual and LOEO meta-analyses approaches, we were able to show that the core regions for ‘Needing’_ST_ (in this case perceiving food when hungry) seems to be the mid-posterior insula.

### ‘Wanting’_ST_ is more of an emotion than Needing’_ST_

Our conjunction results showing consistent activations within the anterior insula for both ‘Wanting’_ST_ and ‘Needing’_ST_ could be related to the fact that this region integrates emotional states and is associated with emotional representation of internal states (Craig, 2010), and increases the significance of external stimuli that are relevant with regard to bodily, affective and sensory information (Young & Nusslock, 2016; Menon and Uddin, 2010). Moreover, based on the fact that the anterior insula plays an important role in awareness (Craig, 2011), our findings suggest that this common activation could be related to our ability to be aware of our wants and needs. It is important to note that the anterior insula was found in the contrast ‘Wanting’_ST_ - ‘Needing’_ST_, but not in ‘Needing’_ST_ - ‘Wanting’_ST_. ‘Wanting’_ST_, viewed as reward seeking is often considered as an emotional state (Panksepp, 2004) that may recruit the anterior insula during reward anticipation without necessity of ‘Needing’_ST_ (Knutson et. al, 2001; see Craig, 2010). Thus, ‘Wanting’_ST_ can be thought of as a form of emotional reaction triggered by a Pavlovian cue that predicts a reward. Whereas ‘Needing’_ST_ (physiological in this case) is usually considered an homeostatic emotion or sometimes a simple sensation, because physiological needs such as hunger do not seem to meet the criteria to be classified as emotions (see Panksepp, 2004), even though they can be seen as homeostatic emotions (Craig, 2003). In light of this, we speculate that, in humans, ‘Wanting’_ST_ can have more emotional power than ‘Needing’_ST_ because of the more consistent recruitment of the anterior insula. Though the anterior insula might contribute to turning ‘Wanting’_ST_ into a more conscious desire/craving (Naqvi et al., 2014; Garavn, 2010), ‘Wanting’_ST_ can also influence behaviour without explicit awareness (Berridge and Robinson, 2003; Strack and Deutsch, 2004; Wei et al., 2017).

### Short- term value for ‘Wanting’_ST_ vs long- term value for ‘Needing’_ST_

While consistent activations were found for both states within the striatum, each seems to recruit a different sub-region with the the ventral and rostral parts, i.e. NAcc, ventromedial caudate and rostroventral putamen more consistenly found for ‘Wanting’_ST_ and the caudo-ventral part of the putamen (that often included the tail of the caudate) more consistently found for ‘Needing’_ST_. This spatial difference could be related to the functional roles of these sub-regions including the coding of short vs. long term values of stimuli. Indeed, our results for ‘Wanting’_ST_ are in line with findings that suggest that ventral striatum is more responsive to reward or its prediction than the dorsal striatum (Schultz et al., 2000) and that rostral striatum, mainly the caudate head, encodes short term or flexible value (Kim and Hikosaka, 2015). This is also in line with the view of ‘Wanting’_ST_ as a moment to moment modulation of a cue that predicts reward in synergy with dopaminergic states (Zhang et al., 2009). In contrast, ‘Needing’_ST_, was more associated with consistent activation within the caudo-ventral putamen (called “putamen tail”, see Kunimatsu et al., 2019) and (to a lesser extent) the caudate tail, both referred to as striatum tail (Amita et al., 2018), regions that acquire long-term values of stimuli based on the historical experience of reward, but not on prediction of rewards (Kunimatsu, et. al., 2019). Thus, in line with theories and previous studies (Kim and Hikosaka, 2013, Zhang et al., 2009; Amita et al., 2018; Kunimatsu et al., 2019), our results might be interpreted as showing that value representation of a wanted vs. needed stimuli rely on distinct regions of the striatum and that this difference could be driven by the temporal aspects or requirement of value processing for each state.

### Directional and activational effect of value

The value assigned to stimuli can have a directional effect or activational effect. The directional effect is linked to choice (preference or action selection) and directs towards or away from stimuli, while the activational effect is related to action initiation, maintenance, and vigor (see Salamone et al., 2018). ‘Wanting’_ST_ AND ‘Needing’_ST_ meta-analytic conjunction showed that both states consistently activate the caudate nucleus (head and body) and anterior insula (discussed above), regions implicated in action selection (Hollon et al., 2014; ito and Doya, 2015; Petzschner et al., 2021) and emotional representation of internal states (Craig, 2010), respectively. The caudate is involved in goal directed behavior (Balleine and O’Doherty, 2010; Knutson & Cooper, 2005), and in the pairing between an action and the value of its consequence (Schwabe & Wolf, 2010), such as on the current state of the organism (see Balleine, 1992). Thus, the caudate is implicated in choice/action selection related value (Hollon et al., 2014; ito and Doya, 2015), and is involved in directional value (Salamone et. al 2016). The implication of the caudate in ‘Wanting’_ST_ AND ‘Needing’_ST_ conjunction suggests both states can influence choice/action selection, i.e., directional value of stimuli. Thus, by doing a meta-analytic conjunction of ‘Wanting’_ST_ AND ‘Needing’_ST_, we were able to show that both ‘Wanting’_ST_ and ‘Needing’_ST_ can influence the directional value of stimuli. However, as we will see, each state seems to rely on distinct neural substrates to compute this directional value.

Directional value for ‘Wanting’_ST_ seems to arise from activity within the dopaminergic system. The VTA and SN, which contain the main dopaminergic neurons were shown to be more consistently activated for ‘Wanting’_ST_ than ‘Needing’_ST_. Our results also show that the regions of the ventral striatum, i.e. the NAcc and the ventromedial caudate and rostroventral putamen (Haber & Knutson, 2010) were more consistently activated for ‘Wanting’_ST_ - ‘Needing’_ST_; as well as the globus pallidus and the ventral pallidum (VP) (not shown). Indeed, incentive salience ‘Wanting’_ST_ is generated when a reward cue is synergistically mixed with the state of mesocorticolimbic circuits (which mainly implicates the VTA, NAcc and pallidum) (Warlow & Berridge, 2021; Zhang et al. 2009). Based on our results, we suggest that the directional value of ‘Wanting’_ST_ towards stimuli comes from the cortico-striato-midbrain pathway, and first starts with the VTA which computes the prediction error that signals change in expected reward prediction (Schultz et al., 1997) and project mesolimbic dopamine to the ventral striatum (NAcc and VP) (Haber & Knutson, 2010). Second, the activity of the NAcc shell which corresponds to ventrolateral putamen in humans is the final path to the directional value of ‘Wanting’_ST_ (Holmes et al., 2010); and it is known that mesolimbic dopamine activation within the NAcc or ventral striatum has strong influence on the dorsal striatum (Tricomi et al., 2009).

As mentioned before, with regard to ‘Needing’_ST_, (the reaction to a needed stimulus); the middle insula, which was found as peak in all of our analyses including our contrasts in favour of ‘Needing’_ST_, seems to be the core regions for ‘Needing’_ST_ (or in this case: when one perceives food while hungry). In this sense, our results confirm that within the insula, it is the middle insula that pairs internal states to relevant external stimuli as argued by Craig (2010). Moreover, our findings dovetail those in the literature that show that the insula plays a role in an “as – if” representation of the bodily state (Damasio, 1994; Naqvi & Bechara, 2010), and that the insula encodes the incentive value of outcomes as a form of incentive memory (Balleine and Dickinson, 2000). Indeed, when hungry or thirsty, the mid-posterior insula simulates future satiety state in the presence of food or water cues for both humans and animals (Chen et al., 2016; Livneh et al., 2020)). Those cues create an interoceptive prediction error (see Barret Simmon, 2015). Based on interoceptive prediction error from mid-posterior insula, the visceromotor cortices (ACC, OFC, anterior insula) make predictions about desired internal states (Barrett and Simmon, 2015), and enhance the value of stimuli and actions that fulfill the predictions (Petzschner et al., 2021). Based on our results, we suggest that the mid-posterior insula prediction error might be the origin of the directional value of ‘Needing’_ST_ in the same logic the VTA does for ‘Wanting’_ST_, i.e. by computing a sort of prediction error that influences cue selection (see Arsenaut et al., 2014); and in this case (i.e. for ‘Needing’_ST_) it’s an interoceptive prediction error (Barrett and Simmon, 2015). In this regard, the directional value of ‘Wanting’_ST_ and that of ‘Needing’_ST_ depend on two different prediction errors: for ‘Wanting’_ST_ the prediction error is computed within the VTA, and for ‘Needing’_ST_ the (interoceptive) prediction error is computed within the mid-posterior insula. Importantly, although we focussed our meta-analysis on the hunger state and the processing of food stimuli, we think that our results can be generalized to other types of needing states and stimuli. Indeed, it is known that the mid-posterior insula receives multimodal sensory interoceptive signals to compute a prediction error (Gerlach et al., 2019).

If both states can give rise to directional/action selection value (albeit differently), only ‘Wanting’_ST_ seems associated to activiational value. Indeed, consistent activations within the NAcc (and ventral striatum) was only found in our ‘Wanting’_ST_ meta-analysis, ‘Wanting’_ST_ - ‘Needing’_ST_ meta-analytic contrast, and even when stringent LOEO analyses were used. ‘Wanting’_ST_ has more (compared to ‘Needing’_ST_) control on activational value because the prediction error signal is sent to the NAcc which (makes those predictions and) has strong influence to the pallidum which has a lot of impact on invigoration of motor action possibly through a more direct connection to the thalamus (Balleine & O’Doherty, 2010; Haber & Knutson, 2010). In line with literature, our results point out that the activational aspects of cue induced ‘Wanting’_ST_ is likely mediated by the mesolimbic dopamine that implicates activation within the central NAcc (see Holmes, 2010; Salamone et al., 2016; Salamone et al., 2018; Salamone et al. , 1997 ; Salamone and Correa, 2002) or ventral striatal regions in general (Haber & Knutson, 2010) which have strong influence on the dorsal striatum (Tricomi et al., 2009). Our results showing that ‘Needing’_ST_ did not consistently activate dopaminergic regions, are in line with the now admitted fact that needs by themselves don’t have activational value (see Salamone et al., 2018) and are not the main source of motivated behaviour (Bindra, 1974; Berridge, 2004), although they can amplify it (Toates, 1994). The fact that ‘Needing’_ST_ has only the directional part (choice/preference or action selection), not the activational one, means that a needed stimulus must still become ‘wanted’, by altering mesolimbic dopamine reactivity and encountering a relevant reward predicting cue (Zhang et al., 2009), in order to have full motivational value (Bindra, 1974; Toates, 1994; Berridge, 2004). Thus, motivation is better explained by incentive salience ‘Wanting’_ST_ than by ‘Needing’_ST_ (Bindra, 1974; Berridge, 2004). Nevertheless, ‘Needing’_ST_, can affect ‘liking’ (see Berridge, 2009) (whether for hunger and food or thirst and water) (Dayan & Balleine, 2002; Balleine 1992), and can create expectation of ‘liking’ through “cognitive desire” (see Berridge, 2012) towards a needed stimulus. This latter is more goal-oriented, and based on declarative memories and on cognitive expectations of act-outcome relations (Berridge, 2012). Thus, ‘Needing’_ST_ generates cognitive desire, but not necessarily ‘Wanting’_ST_ (incentive salience) (Berridge, 2012).

### Conclusion

Our goal was to compare the brain representation of ‘Wanting’_ST_ and ‘Needing’_ST_ related values --two states that guide value attribution and our consumption behaviors. Our results suggest distinct brain systems for both states, with the mesolimbic dopaminergic circuitry as the core for ‘Wanting’_ST_. and the posterior-middle insula for ‘Needing’_ST_. Whereas ‘Needing’_ST_ only provides directional value (through an interoceptive prediction error), ‘Wanting’_ST_ which involves dopamine provides both directional (through a reward prediction error) and activational value (through mesolimbic dopamine in ventral striatum and pallidum). Because ‘Needing’_ST_ does not provide activational value to stimuli, full motivation (directional and activational) to consume depends more on ‘Wanting’_ST_ than on ‘Needing’_ST_, and means that ‘Wanting’_ST_ has more power to activate behavior. This might explain why we consume what we want beyond what we need (Stearn, 2006).

### Limits

We would like to point out some limits to our work. First, ‘Wanting’_ST_ experiments did not include physiologically related stimuli such as food or water, We argue that the experiments for ‘Wanting’_ST_ really expressed ‘wanting’ (see Berridge, 2004) because the contrasts we used focussed on the processing of the cues, not the outcome; and the cues in those experiments triggered the decision for reward seeking. Indeed, ‘wanting’ has been related to decision utility, i.e. the “choice to pursue or consume an outcome” (Berridge & O’Doherty, 2014), induced by a cue (Berridge & Aldridge, 2009). So, although we could not know the mesolimbic state of participants in those experiments, the behavioral situations, i.e. a cue that triggered a decision to seek reward, do seem to induce ‘wanting’ (see Berridge, 2004). Furthermore, ‘Wanting’_ST_ or motivation activates a general system, regardless of the type of stimulus (Bouton, 2016). Thus, though most stimuli for ‘Wanting’_ST_ were money or points, the mechanism for cue induced decision for reward seeking, i.e. ‘Wanting’_ST_ is the same.

Second, ‘Needing’_ST_ contrats only focused on hunger and the processing of food (physiologically related stimuli). Though our method could seem dependent on the type of physiological state, i.e. hunger, and thus the type of needed stimuli, i.e. food; our findings and interpretation seem to go in the same direction than studies and theories that suggest an integrative role of all physiological states within the insula. In that sense Gerlach et al. (2020) results suggest that dense multimodal sensory prediction errors converge in the posterior insula to guide interoception. Based on that, it has been argued that that the insula represents physiological state in cue-independent spontaneous activity, which is then modified directionally by cues that predict water or food availability (Namboodiri & Stuber, 2020). This suggests that our findings regarding the activation of mid-posterior insula might not be hunger/food dependent but rather be resulting from a more general role of the insula with regard to bodily states, specifically serving interoceptive inference (Allen, 2020). Though, there are justifications for our methodology in general, it would still be interesting for future studies to test the difference between ‘Wanting’_ST_ and ‘Needing’_ST_ with more control on the dopaminergic state, on the type of physiological states and on the type of stimuli.

Nevertheless, wanting and/or needing go way beyond eating, drinking or winning money or points. The distinction between wanting vs needing can apply to virtually any decision, and can even be viewed as philosophical or phenomenological concepts. In that sense, they go way beyond our study and the tools we have used. Thus, our study should be viewed as testing some manifestation of wanting and/or needing rather than testing the general phenomena.

## ACKNOWLEDGMENTS

The research was supported in part by NSERC Discovery Grant #RGPIN-2018-05698, UdeM institutional funds, and Mitacs Grant #IT20458.

## AUTHOR CONTRIBUTION

**Juvénal Bosulu**: Designed the study, performed the database search, performed data analysis, interpretation, and wrote the manuscript. **Sébastien Hétu**: Designed the study, revised the manuscript and provided critical feedbacks. **Max-Antoine Allaire**: Performed the database search, revised the manuscript and provided critical feedbacks. **Laurence Tremblay-Grenier**: Performed the database search, revised the manuscript and provided critical feedbacks. **Yi Luo**: Revised the manuscript and provided critical feedbacks. **Simon Eichkoff**: Revised the manuscript and provided critical feedbacks. All authors contributed to and approved the final manuscript version.

## DATA AVAILABILITY STATEMENT

All Data are available upon request.

