## supplementary material for "‘Wanting’ versus ‘Needing’ related value: an fMRI meta-analysis"

### PRISMA for Wanting$_{ST}$

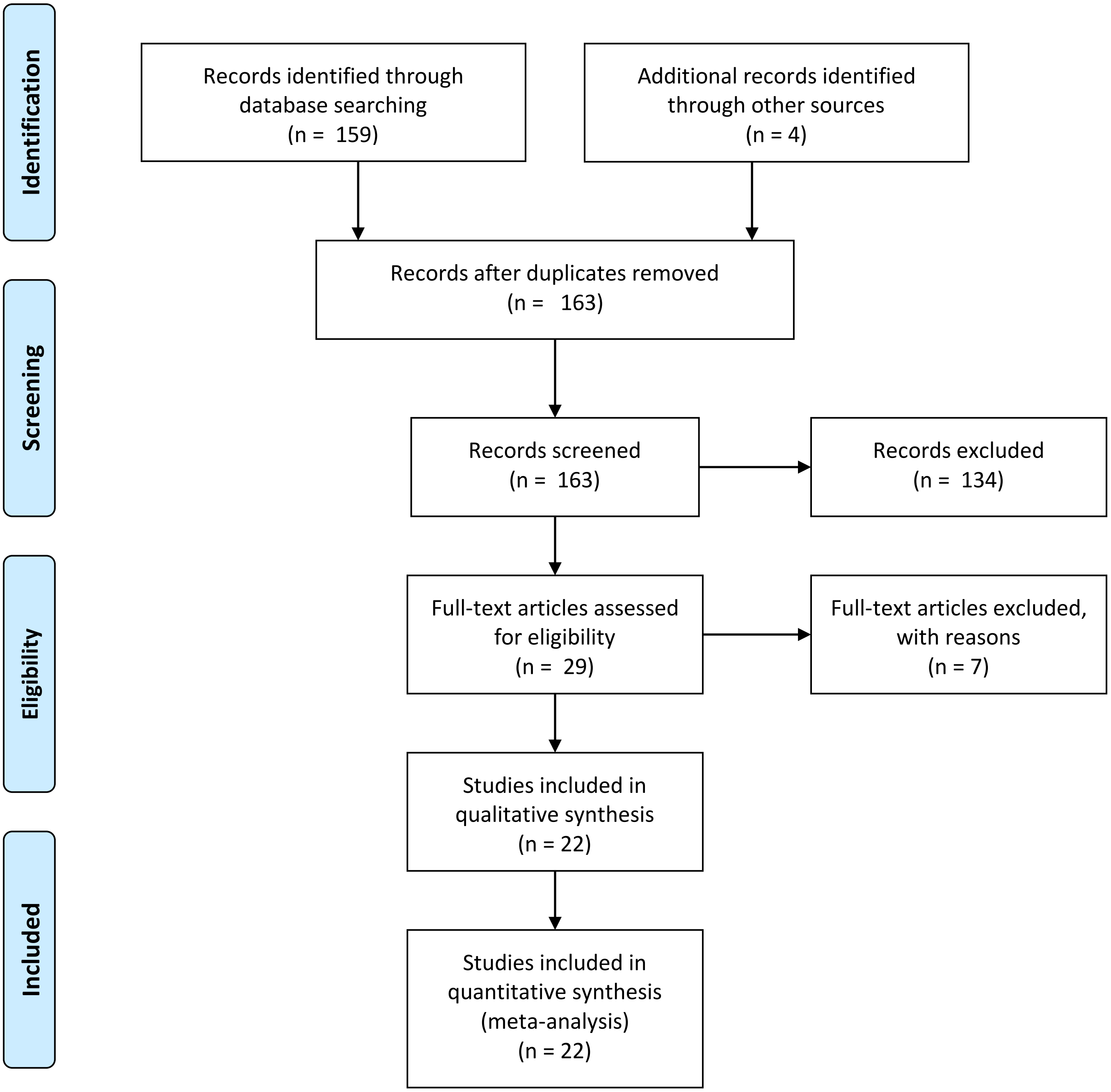

**PRISMA Needing$_{ST}$**

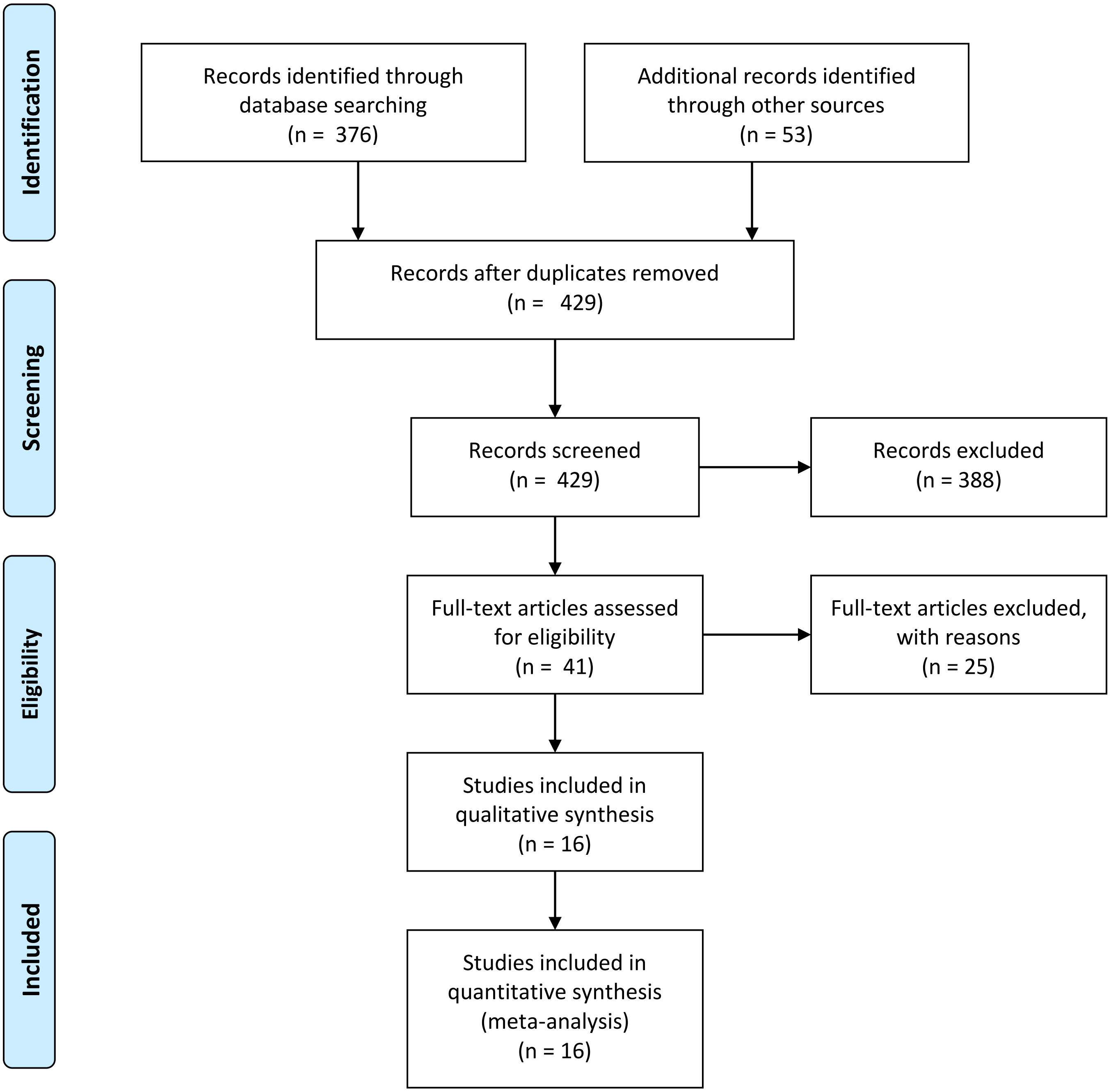

### Leave one experiment out (LOEO) analysis

**WANTING$_{ST}$**

Threshold
-- p value = 1
-- intensity = -1.633123935319537e+16
-- cluster size = 5
Number of clusters found: 10
---------------------
Cluster 1
Number of voxels: 354
Peak MNI coordinate: 8 -26 -22
Peak MNI coordinate region: // Right Brainstem // Midbrain // undefined // undefined // undefined // undefined
Peak intensity: 1
### voxels structure

| | |
|---|---|
| 354 | --TOTAL # VOXELS-- |
| 354 | Midbrain |
| 199 | Left Brainstem |
| 155 | Right Brainstem |
| 12 | Gray Matter |
| 6 | Substania Nigra |
| 6 | Red Nucleus |

---------------------
Cluster 2
Number of voxels: 527
Peak MNI coordinate: 18 4 -12
Peak MNI coordinate region: // Right Cerebrum // Sub-lobar // Lentiform Nucleus // Gray Matter // Putamen // undefined
Peak intensity: 1
### voxels structure

| | |
|---|---|
| 527 | --TOTAL # VOXELS-- |
| 527 | Right Cerebrum |
| 523 | Sub-lobar |
| 443 | Gray Matter |
| 244 | Lentiform Nucleus |
| 210 | Putamen |
| 204 | Caudate_R (aal) |
| 171 | Caudate |
| 131 | Putamen_R (aal) |
| 126 | Caudate Head |
| 81 | White Matter |
| 77 | Extra-Nuclear |
| 77 | Pallidum_R (aal) |
| 45 | Caudate Body |
| 31 | Lateral Globus Pallidus |
| 3 | Medial Globus Pallidus |
| 3 | Frontal Lobe |
| 3 | Subcallosal Gyrus |
| 3 | Lateral Ventricle |
| 3 | Cerebro-Spinal Fluid |
| 1 | Limbic Lobe |
| 1 | Anterior Cingulate |

---------------------

Cluster 3
Number of voxels: 480
Peak MNI coordinate: -4 10 -12
Peak MNI coordinate region: // Left Cerebrum // Frontal Lobe // Subcallosal Gyrus // Gray Matter // brodmann area 25 // Caudate_L (aal)
Peak intensity: 1
### voxels structure

| | |
|---|---|
| 480 | --TOTAL # VOXELS-- |
| 480 | Left Cerebrum |
| 455 | Sub-lobar |
| 336 | Gray Matter |
| 195 | Lentiform Nucleus |
| 184 | Putamen_L (aal) |
| 180 | Caudate_L (aal) |
| 177 | Putamen |
| 144 | White Matter |
| 125 | Extra-Nuclear |
| 110 | Caudate |
| 84 | Caudate Head |
| 45 | Pallidum_L (aal) |
| 26 | Caudate Body |
| 16 | Lateral Globus Pallidus |
| 14 | Frontal Lobe |
| 14 | Subcallosal Gyrus |
| 11 | Limbic Lobe |
| 10 | Anterior Cingulate |
| 3 | brodmann area 25 |
| 2 | brodmann area 13 |
| 1 | Medial Globus Pallidus |

----------------------
Cluster 4
Number of voxels: 173
Peak MNI coordinate: 32 22 -6
Peak MNI coordinate region: // Right Cerebrum // Frontal Lobe // Inferior Frontal Gyrus // White Matter // undefined // undefined
Peak intensity: 1
### voxels structure

| | |
|---|---|
| 173 | --TOTAL # VOXELS-- |
| 173 | Right Cerebrum |
| 112 | Insula_R (aal) |
| 105 | White Matter |
| 105 | Sub-lobar |
| 68 | Frontal Lobe |
| 60 | Inferior Frontal Gyrus |
| 60 | Gray Matter |
| 59 | Insula |
| 40 | Extra-Nuclear |
| 24 | brodmann area 13 |
| 19 | brodmann area 47 |
| 11 | Sub-Gyral |
| 8 | brodmann area 45 |
| 5 | Putamen_R (aal) |
| 3 | Claustrum |
| 1 | Frontal_Inf_Tri_R (aal) |

----------------------
Cluster 5

Number of voxels: 174
Peak MNI coordinate: -32 16 -2
Peak MNI coordinate region: // Left Cerebrum // Sub-lobar // Insula // Gray Matter // brodmann area 47 // Insula_L (aal)
Peak intensity: 1

| # voxels | structure |
|---|---|
| 174 | --TOTAL # VOXELS-- |
| 174 | Left Cerebrum |
| 169 | Sub-lobar |
| 143 | Insula_L (aal) |
| 107 | White Matter |
| 98 | Insula |
| 60 | Gray Matter |
| 52 | Extra-Nuclear |
| 31 | brodmann area 13 |
| 17 | Claustrum |
| 7 | brodmann area 47 |
| 7 | Inferior Frontal Gyrus |
| 5 | Frontal Lobe |
| 4 | Frontal_Inf_Tri_L (aal) |
| 3 | Putamen_L (aal) |
| 2 | brodmann area 45 |

----------------------

Cluster 6
Number of voxels: 98
Peak MNI coordinate: 38 34 24
Peak MNI coordinate region: // Right Cerebrum // Frontal Lobe // Middle Frontal Gyrus // White Matter // undefined // Frontal_Inf_Tri_R (aal)
Peak intensity: 1

| # voxels | structure |
|---|---|
| 98 | --TOTAL # VOXELS-- |
| 98 | Frontal Lobe |
| 98 | Right Cerebrum |
| 95 | White Matter |
| 86 | Middle Frontal Gyrus |
| 85 | Frontal_Mid_R (aal) |
| 13 | Frontal_Inf_Tri_R (aal) |
| 10 | Sub-Gyral |
| 3 | brodmann area 9 |
| 3 | Gray Matter |
| 2 | Superior Frontal Gyrus |

----------------------

Cluster 7
Number of voxels: 50
Peak MNI coordinate: -44 -38 42
Peak MNI coordinate region: // Left Cerebrum // Parietal Lobe // Inferior Parietal Lobule // White Matter // undefined // Parietal_Inf_L (aal)
Peak intensity: 1

| # voxels | structure |
|---|---|
| 50 | --TOTAL # VOXELS-- |
| 50 | Left Cerebrum |
| 50 | Parietal Lobe |
| 38 | Inferior Parietal Lobule |
| 27 | Postcentral_L (aal) |
| 23 | Gray Matter |
| 23 | White Matter |

23 Parietal_Inf_L (aal)
22 brodmann area 40
8 Postcentral Gyrus
4 Sub-Gyral
1 brodmann area 2

----------------------

Cluster 8
Number of voxels: 163
Peak MNI coordinate: -4 8 46
Peak MNI coordinate region: // Left Cerebrum // Limbic Lobe // Cingulate Gyrus // Gray Matter // brodmann area 32 // Supp_Motor_Area_L (aal)
Peak intensity: 1
### voxels structure
163 --TOTAL # VOXELS--
148 Frontal Lobe
143 Supp_Motor_Area_L (aal)
134 Left Cerebrum
96 Medial Frontal Gyrus
82 Gray Matter
52 Superior Frontal Gyrus
49 White Matter
46 brodmann area 6
33 brodmann area 32
24 Right Cerebrum
18 Supp_Motor_Area_R (aal)
10 Limbic Lobe
10 Cingulate Gyrus
5 Inter-Hemispheric
2 brodmann area 24
1 brodmann area 8

----------------------

Cluster 9
Number of voxels: 65
Peak MNI coordinate: 28 -6 48
Peak MNI coordinate region: // Right Cerebrum // Frontal Lobe // Middle Frontal Gyrus // White Matter // undefined // Precentral_R (aal)
Peak intensity: 1
### voxels structure
65 --TOTAL # VOXELS--
65 Frontal Lobe
65 Right Cerebrum
58 Middle Frontal Gyrus
40 White Matter
25 brodmann area 6
25 Gray Matter
21 Frontal_Mid_R (aal)
20 Precentral_R (aal)
11 Frontal_Sup_R (aal)
7 Sub-Gyral

----------------------

Cluster 10
Number of voxels: 78
Peak MNI coordinate: -28 -6 52
Peak MNI coordinate region: // Left Cerebrum // Frontal Lobe // Middle Frontal Gyrus // White Matter // undefined // Frontal_Mid_L (aal)
Peak intensity: 1

| # voxels | structure |
|---|---|
| 78 | --TOTAL # VOXELS-- |
| 78 | Frontal Lobe |
| 78 | Left Cerebrum |
| 62 | Middle Frontal Gyrus |
| 55 | Precentral_L (aal) |
| 39 | brodmann area 6 |
| 39 | Gray Matter |
| 34 | White Matter |
| 16 | Frontal_Sup_L (aal) |
| 12 | Precentral Gyrus |
| 7 | Frontal_Mid_L (aal) |
| 4 | Sub-Gyral |

>>

**NEEDING$_{ST}$**

Threshold
-- p value = 1
-- intensity = -1.633123935319537e+16
-- cluster size = 5
Number of clusters found: 3
----------------------
Cluster 1
Number of voxels: 8
Peak MNI coordinate: 42 4 -12
Peak MNI coordinate region: // Right Cerebrum // Sub-lobar // Extra-Nuclear // Gray Matter // brodmann area 13 // Insula_R (aal)
Peak intensity: 1

| # voxels | structure |
|---|---|
| 8 | --TOTAL # VOXELS-- |
| 8 | Insula_R (aal) |
| 8 | Right Cerebrum |
| 5 | Sub-lobar |
| 5 | Extra-Nuclear |
| 4 | brodmann area 13 |
| 4 | Gray Matter |
| 3 | Sub-Gyral |
| 3 | Temporal Lobe |
| 2 | White Matter |

----------------------
Cluster 2
Number of voxels: 92
Peak MNI coordinate: -36 -2 -10
Peak MNI coordinate region: // Left Cerebrum // Sub-lobar // Extra-Nuclear // White Matter // undefined // Insula_L (aal)
Peak intensity: 1

| # voxels | structure |
|---|---|
| 92 | --TOTAL # VOXELS-- |
| 92 | Left Cerebrum |
| 89 | Sub-lobar |
| 85 | White Matter |
| 77 | Insula |
| 71 | Insula_L (aal) |
| 9 | Extra-Nuclear |

7 Gray Matter
4 brodmann area 13
3 Sub-Gyral
3 Claustrum
3 Temporal Lobe

----------------------

Cluster 3
Number of voxels: 39
Peak MNI coordinate: 42 -6 -2
Peak MNI coordinate region: // Right Cerebrum // Sub-lobar // Insula // White Matter // undefined // Insula_R (aal)
Peak intensity: 1

| # voxels | structure |
|---|---|
| 39 | --TOTAL # VOXELS-- |
| 39 | Sub-lobar |
| 39 | Right Cerebrum |
| 22 | White Matter |
| 18 | Extra-Nuclear |
| 18 | Insula_R (aal) |
| 17 | Gray Matter |
| 14 | Insula |
| 12 | Putamen_R (aal) |
| 8 | brodmann area 13 |
| 7 | Claustrum |

>>
